# ICOS limits memory-like properties and function of exhausted PD-1^+^ CD8 T cells

**DOI:** 10.1101/2024.09.16.611518

**Authors:** Etienne Humblin, Isabel Korpas, Nataliya Prokhnevska, Abishek Vaidya, Jiahua Lu, Verena van der Heide, Dan Filipescu, Tesia Bobrowski, Adam Marks, Matthew D. Park, Emily Bernstein, Brian D Brown, Amaia Lujambio, David Dominguez-Sola, Brad R Rosenberg, Alice O. Kamphorst

**Affiliations:** Marc and Jennifer Lipschultz Precision Immunology Institute, Icahn School of Medicine at Mount Sinai, New York, NY, USA; Department of Immunology and Immunotherapy, ISMMS, New York, NY, USA; Tisch Cancer Institute, ISMMS, New York, NY, USA; Department of Genetics and Genomics, ISMMS, New York, NY, USA; Department of Oncological Sciences, ISMMS, New York, NY, USA; Department of Microbiology, ISMMS, New York, NY, USA

**Keywords:** ICOS, costimulation, exhaustion, FoxO1, CD8 T cells, PD-1, T cell differentiation, progenitor exhausted

## Abstract

During persistent antigen stimulation, PD-1^+^CD8 T cells are maintained by progenitor exhausted PD-1^+^TCF-1^+^CD8 T cells (Tpex). Tpex respond to PD-1 blockade, and regulation of Tpex differentiation into more functional Tex is of major interest for cancer immunotherapies. Tpex express high levels of Inducible Costimulator (ICOS), but the role of ICOS for PD-1^+^CD8 T cell responses has not been addressed. In chronic infection, ICOS-deficiency increased both number and quality of virus-specific CD8 T cells, with accumulation of effector-like Tex due to enhanced survival. Mechanistically, loss of ICOS signaling potentiated FoxO1 activity and memory-like features of Tpex. In mice with established chronic infection, ICOS-Ligand blockade resulted in expansion of effector-like Tex and reduction in viral load. In a mouse model of hepatocellular carcinoma, ICOS inhibition improved cytokine production by tumor-specific PD-1^+^CD8 T cells and delayed tumor growth. Overall, we show that ICOS limits CD8 T cell responses during chronic antigen exposure.

## INTRODUCTION

Antigen persistence, as in chronic viral infection or cancer, leads to the differentiation of hypofunctional exhausted Programmed Cell Death-1 (PD-1)^+^ CD8 T cells^1,2^. T Cell Factor-1 (TCF-1)^+^ PD-1^+^ CD8 T cells act as progenitors (Tpex)^3–6^, that give rise to differentiated exhausted cells (Tex), which can retain some effector function (effector-like Tex), or become dysfunctional terminally exhausted cells (Terminal Tex)^7,8^. Tpex are critical for sustaining PD-1^+^ CD8 T cell responses and provide the proliferative burst in response to PD-1-targeted therapy^3,9,10^. Therefore, there is great interest to understand the signals and molecular programs that control Tpex differentiation.

Intratumoral Tpex can be found in antigen-presenting cell (APC)-rich niches^11,12^. We recently showed in patients with hepatocellular carcinoma (HCC), that Tpex colocalization with CD4 T cells and mature dendritic cells (DC) was associated with expansion of PD-1^+^ effector-like CD8 T cells and response to PD-1-targeted therapy^12^. APC-rich niches and CD4-help may provide critical signals, through costimulation or cytokines, to guide effective Tpex differentiation and survival^8,13–16^. Tpex express high levels of costimulatory molecules, such as CD28 and Inducible Costimulator (ICOS) ^3^. We previously found that CD28 costimulation was essential for the success of PD-1-targeted therapy^17^ and we recently showed that Tpex continuously require low-level CD28 signaling to survive and rely on stronger CD28 costimulation to differentiate^18^, supporting the key role of costimulation for differentiation of PD-1^+^ CD8 T cells.

Tpex share similarities with memory CD8 T cells (Tmem) as well as follicular helper CD4 T cells (Tfh)^3^, but the requirement for specific gene programs for Tpex function and differentiation remains to be better defined. ICOS is induced upon TCR signaling and binds to B7 family member ICOS-Ligand (ICOSL/B7-H2), which is constitutively expressed on APCs and non-lymphoid cells, including tumor and epithelial cells^19–23^. Notably, ICOS signaling is necessary to induce Tfh differentiation and maintenance^24–26^, through forkhead box O1 (FoxO1) inactivation^26^. However, FoxO1 is a master positive regulator of memory T cell development^27–30^ and in chronic viral infection, FoxO1 is indispensable for Tpex survival and long-term maintenance of the exhausted CD8 T cell population^31,32^. However, the relationship between ICOS signaling and FoxO1 regulation on CD8 T cells has not been established.

Peng et al. have shown that ICOS is dispensable for the effector CD8 T cell response and differentiation of circulating memory cells following acute lymphocytic choriomeningitis virus (LCMV) infection. However, ICOS signaling was necessary for the development and maintenance of tissue resident memory CD8 T cells^22^. Here, using LCMV chronic infection and a murine model of liver cancer, we found that ICOS deficiency improves PD-1^+^ CD8 T cell responses. Mechanistically, we uncovered that FoxO1 activity is enhanced in the absence of ICOS. ICOS deficiency on CD8 T cells or therapeutic blockade of ICOS-Ligand improved memory-like features of Tpex, as well as survival and functionality of effector-like Tex. Overall, we demonstrate that sustained ICOS costimulation is detrimental to PD-1^+^ CD8 T cells during persistent antigen exposure, highlighting the contrasting effects of different costimulatory receptors on the regulation of T cell responses.

## RESULTS

### ICOS deficiency promotes expansion of effector-like PD-1^+^ CD8 T cells

To address the role of the costimulatory molecule ICOS on PD-1^+^ CD8 T cells during chronic antigen stimulation, we first used a CRISPR-Cas9 approach to delete *Icos* on virus-specific CD8 T cells. Naïve transgenic P14 (CD8 T cells specific for LCMV-GP33) were electroporated with Cas9-ribonucleoprotein complex targeting *Icos* (sgIcos) or a control gene not expressed in T cells (sgCtrl) (**Fig.S1A**)^33,34^. Transfected P14 were transferred into naïve recipient mice before infection with LCMV clone 13 (**Fig.1A)**. We compared the kinetics of P14 responses in the blood and confirmed efficiency of ICOS deletion (**Fig. S1B**). 8 days post infection (DPI), there was similar expansion of sgCtrl and sgIcos P14, suggesting that ICOS signaling is not required for priming and early expansion of virus-specific CD8 T cells following LCMV clone 13 infection (**Fig. 1B**). Following initial expansion, ICOS-deficient P14 underwent slower contraction compared to control P14, resulting in a significantly higher frequency of circulating ICOS-deficient P14 during the chronic phase of infection (**Fig. 1B**). Accordingly, in mice with established chronic infection, we observed higher frequency and numbers of ICOS-deficient P14 compared to control in spleen and lung (**Fig.1C** and **Fig.S1C-D)**.

**Fig. 1:**
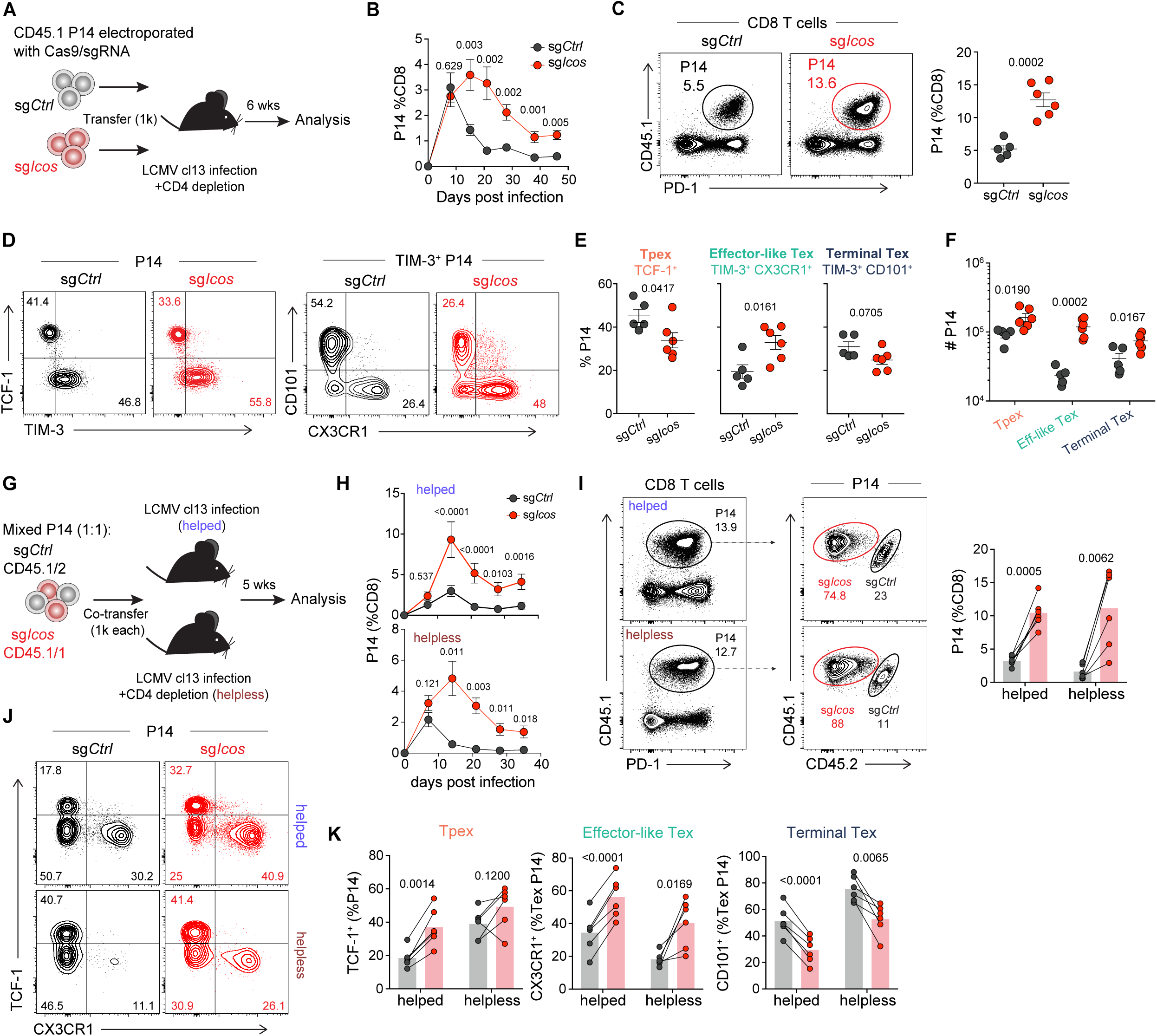
ICOS-deficiency promotes expansion of virus-specific CD8 T cells in LCMV chronic infection. **(A)** Experimental layout for (B-F): CD45.1 P14 were electroporated with Cas9/single guide RNA Control (sgCtrl, black) or Cas9/sgRNA targeting Icos (sgIcos, red), and transferred (1,000 cells) into recipient C57BL/6J mice (depleted of CD4 T cells) the day before LCMV clone 13 infection. P14 were analyzed in the spleen after 42 days. **(B)** P14 frequency in blood. **(C)** Representative contour plot and graph showing frequency of P14 in spleen. **(D)** Contour plot showing expression of TCF-1 and TIM3 (left), CX3CR1 and CD101 (right). **(E)** Frequency and **(F)** absolute number of progenitor (TCF-1^+^), effector-like Tex (TIM3^+^ CX3CR1^+^) and terminal Tex (TIM3^+^ CD101^+^) P14 sgCtrl and sgIcos. **(G)** Experimental layout for (H-K): P14 were electroporated with Cas9/sgCtrl or Cas9/sgIcos and co-transferred (1,000 cells each) to C57BL/6J mice (depleted of CD4 T cells=helpless, or not=helped) the day before LCMV clone 13 infection. P14 were analyzed in spleen after 35 days. **(H)** P14 frequency in blood. **(I)** Contour plot and graph showing frequency of splenic co-transferred P14. **(J)** P14 expression of TCF-1 and CX3CR1. **(K)** Frequency of progenitor, effector-like Tex and terminal Tex. Data in (B-F) and (H-K) are representative of 3 independent experiments with five to six mice per group. Error bars show standard error of the mean (SEM). (H-K) Connecting lines between symbols indicate cells analyzed from the same mouse. (B-F) unpaired Student’s t-test; (H-K) paired Student’s t-test.

Next, we examined how loss of ICOS affects the differentiation of chronically stimulated CD8 T cells. ICOS is mainly expressed by TCF-1^+^ Tpex. Lower levels of ICOS are found on about 50% of CX3CR1^+^ TIM3^+^ effector-like cells, and only minimal ICOS expression was detected on CX3CR1^neg^ CD101^+^ TIM3^+^ terminal Tex (**Fig.S1E**). When we analyzed the distribution of ICOS-deficient P14 among different subsets, we observed decreased relative frequency of Tpex and terminal Tex subsets and an increase in the TIM3^+^ CX3CR1^+^ effector-like subset (**Fig.1D-E**). However, the absolute number of all subsets were higher in mice that received ICOS-deficient P14 (**Fig.1F**), indicating that the observed lower frequency of Tpex is due to the higher expansion of effector-like P14. The substantial increase in the absolute number of effector-like P14 suggests that loss of ICOS leads to the preferential expansion of effector-like PD-1^+^ CD8 T cells. Of note, there are very few effector-like PD-1^+^ CD8 T cells in liver^35^, which may explain why we did not find a difference in frequency between sgCtrl and sgIcos P14 in this organ (**Fig.S1C-D)**.

CX3CR1^+^ effector-like PD-1^+^ CD8 T cells represent a transitory population of virus-specific cells, differentiated from TCF-1^+^ progenitors, that preserve some effector function. Effector-like cells are essential for viral control^7,8^, and their differentiation is enhanced by CD4 helper T cells^8,15,16^. We observed an accumulation of effector-like ICOS-deficient P14 in the helpless model (CD4-depleted) of LCMV chronic infection. To determine if our observations can be recapitulated in a helped setting, we co-transferred control and sgIcos P14 into WT or CD4-depleted recipient mice before infection with LCMV clone 13 (**Fig. 1G**). In accordance with our previous observations, ICOS-deficient P14 did not display compromised expansion, and instead, accumulated at higher frequencies than control P14 in both helped and helpless settings (**Fig.1H**). At 35 DPI, ICOS-deficient P14 represented around 80% of P14 in spleen (**Fig.1I**), and there was a substantial increase in effector-like subset in helped and helpless settings (**Fig.1J-K**). There was also a higher frequency of Tpex among ICOS-deficient P14 in the helped LCMV chronic infection model, highlighting that a lack of ICOS signaling is not detrimental to progenitor exhausted CD8 T cells. Together, these data indicate that ICOS signaling restrains PD-1^+^ CD8 T cells regardless of CD4-help.

### ICOS signaling limits FoxO1 activity and Tpex memory-like features

Since ICOS is preferentially expressed by Tpex (**Fig. S1E**), we first sought to address how ICOS signaling regulates Tpex gene expression. We performed transcriptional analysis of control and ICOS-deficient Tpex P14 sorted from helpless LCMV chronically infected mice. We observed 148 genes upregulated in ICOS-deficient Tpex, while 97 genes were downregulated compared to control (**Fig.S2A**). We compared genes upregulated in ICOS-deficient Tpex with publicly available chromatin immunoprecipitation sequencing (ChIP-seq) data and found enrichment of Forkhead box O1 (FoxO1) target genes (**Fig.2A**). Recent work has shown that FoxO1 overexpression in CAR T cells was associated with improved memory features and functionality^36,37^, and identified a FoxO1 regulon as an accurate readout for FoxO1 transcriptional activity^36^. ICOS-deficient Tpex were significantly enriched in the FoxO1 regulon signature (**Fig.2B**), suggesting improved FoxO1 activity in the absence of ICOS signaling. Activation of PI3K/Akt pathway is responsible for FoxO1 protein phosphorylation leading to its translocation into the cytoplasm and proteasomal degradation^38^. Thus, FoxO1 activity is tightly linked to its nuclear retention^31^. To further evaluate FoxO1 activity, we used confocal microcopy and assessed FoxO1 nuclear localization. ICOS-deficient Tpex displayed increased nuclear FoxO1 localization compared to control Tpex (**Fig.2C**). Together these data suggest that lack of ICOS signaling limits FoxO1 cytoplasmic translocation, resulting in increased FoxO1 activity in PD-1^+^ CD8 T cells.

**Fig. 2:**
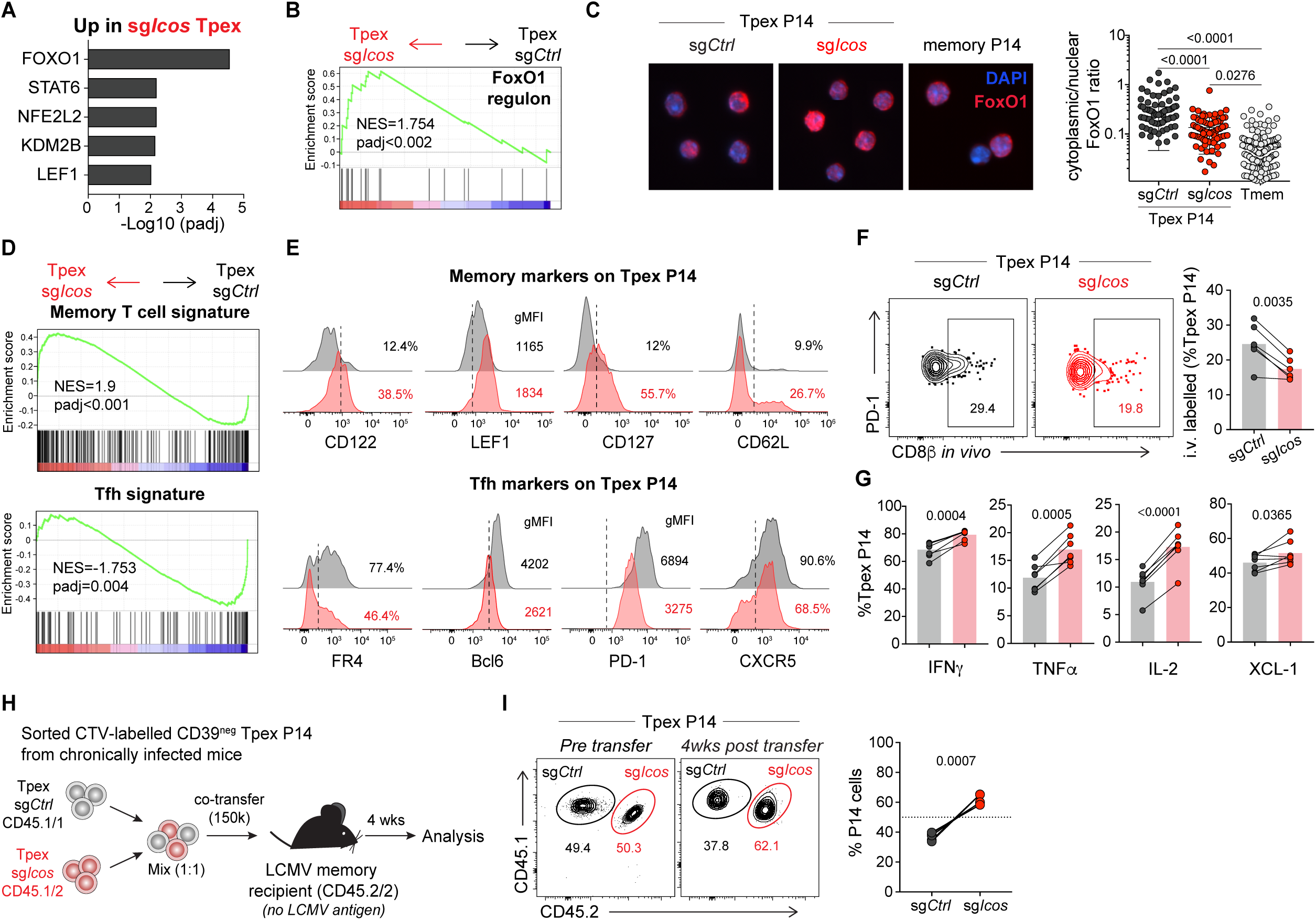
ICOS signaling limits FoxO1 activity and Tpex memory features. (A-C) sgCtrl and sgIcos Tpex (CD39^neg^) P14 were sorted from spleens of LCMV-chronically infected mice, (A-B) RNA was isolated for transcriptional analysis and (C) cells were stained for confocal microscopy. **(A-B)** Gene set enrichment analysis (GSEA) of sgIcos Tpex relative to **(A)** CHEA (ChIP-X Enrichment Analysis, enrichr) or **(B)** FoxO1 regulon gene set (Doan *et al.*, 2023). **(C)** Representative images of FoxO1 (red) localization in Tpex or memory P14, nuclear staining by DAPI (blue); graph shows ratio of FoxO1 Mean Fluorescent Intensity (MFI) in cytoplasm compared to nucleus, symbols represent individual cells. **(D)** GSEA of Memory CD8 T cell and Tfh signatures in ICOS-deficient Tpex compared to control Tpex. **(E)** Representative histograms of proteins associated with memory cells (LEF1, CD62L, CD127, CD122) or Tfh (Bcl6, PD-1, FR4, CXCR5) in Tpex sgCtrl (black) and sgIcos (red) co-transferred P14. Vertical dashed line indicate positivity. **(F)** Mice received intravenous (i.v.) anti-CD8β, contour plots and graphs show frequency of control and sgIcos Tpex P14 susceptible to intravascular staining. **(G)** Frequency of cytokine-producing cells among sgCtrl (black) and sgIcos (red) Tpex P14. **(H)** Experimental layout for (I): Sorted CD39^neg^ Tpex P14 were CTV-labeled and mixed at 1:1 ratio (150k total cells) for co-transfer into B6 recipients that had recovered from LCMV acute infection (memory mice, 65 days post infection). **(I)** Representative contour plots and graph show relative frequencies of sgIcos and sgCtrl Tpex pre and 4 weeks post transfer. Data in (C) are representative from 2 independent experiments with two mice per group. Data in (E and G) are representative of 3 independent experiments with five to six mice per group. Data in (F) are pooled from 2 independent experiments with 3 mice per group. Data in (I) are representative of 2 independent experiments with 4 to 7 mice per groups. (F-I) Connecting lines between symbols indicate cells analyzed from the same mouse, with bars showing the mean value of the group. (C) ANOVA with Sidak’s correction for multiple comparisons; (F-I) paired Student’s t-test

Consistent with enhanced FoxO1 activity, Gene Set Enrichment Analysis (GSEA) showed an increase in memory signature among ICOS-deficient Tpex (**Fig.2D)**. In contrast, we observed that Tfh signature was enriched in control Tpex (**Fig.2D)**. Among genes upregulated in ICOS-deficient Tpex we identified genes involved in memory T cell formation (*Eomes, Lef1, Cd28*), homing (*S1pr1)* and survival (*Il7r)* (**Fig.S2A-B)**. We confirmed these findings by staining ICOS-deficient and control Tpex to assess expression of memory and Tfh-associated proteins. We found that ICOS-deficient Tpex upregulated protein markers associated with memory T cells, such as CD122, LEF1, CD127, CD62L and CD28 (**Fig.2E and S2C**). In contrast, Tfh-associated markers such as FR4, Bcl6, PD-1 and CXCR5 were decreased in ICOS-deficient Tpex (**Fig.2E and S2C**), in accordance with the key role of ICOS and FoxO1 inactivation in Tfh^26^.

Previous studies demonstrated preferential localization of Tpex in the splenic white pulp of LCMV chronically infected animals^3,39,40^. It also have been shown that localization of CD8 T cells in the T cell zone of the white pulp is critical for the development of central memory CD8 T cells^41^. To evaluate potential changes in localization/homing of ICOS-deficient Tpex, we performed in vivo intravascular staining by intravenous (i.v.) injection of anti-CD8β.2 to evaluate cell accessibility to the blood stream (red pulp). The majority of Tpex were not stained by i.v. labelling, with a significantly lower frequency among ICOS-deficient Tpex, indicating increased residence in splenic white pulp in the absence of ICOS signaling (**Fig.2F**). Moreover, after 1:1 co-transfer of control and sgIcos P14 into recipient mice before infection with LCMV clone 13, ICOS-deficient P14 accounted for around 90% of P14 in lymph nodes in established chronic infection (**Fig.S2D**). Also, consistent with their preferential localization in lymphoid tissue, we found a high proportion of Tpex among P14 in lymph nodes

In accordance with improved memory-like features, we observed that ICOS-deficient Tpex exhibited a higher capacity to produce cytokines upon re-stimulation, including increased polyfunctionality as assessed by the number of different cytokines co-produced per cell (**Fig.2G and S2D**). Recent work demonstrated that Tpex can survive in the absence of cognate antigen with partial acquisition of memory T cell properties^42^. To assess if ICOS-deficient Tpex would persist better in the absence of antigen, we isolated control and ICOS-deficient P14 Tpex from LCMV chronically infected mice, and co-transferred (1:1) CTV-labelled sorted Tpex into LCMV memory recipient mice (**Fig.2H**). One month post-transfer, ICOS-deficient P14 represented 65% of transferred cells (**Fig.2I**), demonstrating that ICOS-deficient Tpex have a greater ability to survive without antigen, a key property of memory T cells. Overall, these data suggest that sustained ICOS signaling during antigen persistence limits Tpex memory-like properties.

### ICOS deficiency drives accumulation of effector-like Tex

To gain a more granular understanding of the role of ICOS signaling for differentiation of virus-specific CD8 T cells during antigen persistence, we performed single cell RNA sequencing of control and sgIcos P14 sorted from LCMV chronically infected animals (**Fig.S3A**). We identified five different clusters among P14 (**Fig.3A-B**). We found a Tpex cluster defined by canonical progenitor markers (*Tcf7, Lef1, Bcl6, Xcl1, Il7r, Slamf6*), as well as two clusters of differentiated Tex expressing *Havcr2*, *Gzmb*, *Entpd1*, and higher levels of *Tox* and *Pdcd1*(**Fig.3A-B and S3B**). Tex were either effector-like, with higher expression of genes associated with effector T cell differentiation and migration (*Tbx21, Zeb2, S1pr1, Cx3cr1)* or terminally exhausted, with higher expression of inhibitory receptors (*Cd101, Cd160, Cd200r, Cd244a*)(**Fig.3A-B and S3B)**. We also identified two small clusters of cells defined by higher expression of Interferon Stimulated Genes (ISG) or Heat Shock Protein (HSP) genes (**Fig.3A-B**). While both control and ICOS-deficient P14 were found in every cluster, ICOS-deficient cells were specifically enriched among effector-like Tex, whereas control cells were more represented among terminal Tex (**Fig.3C**).

**Fig. 3:**
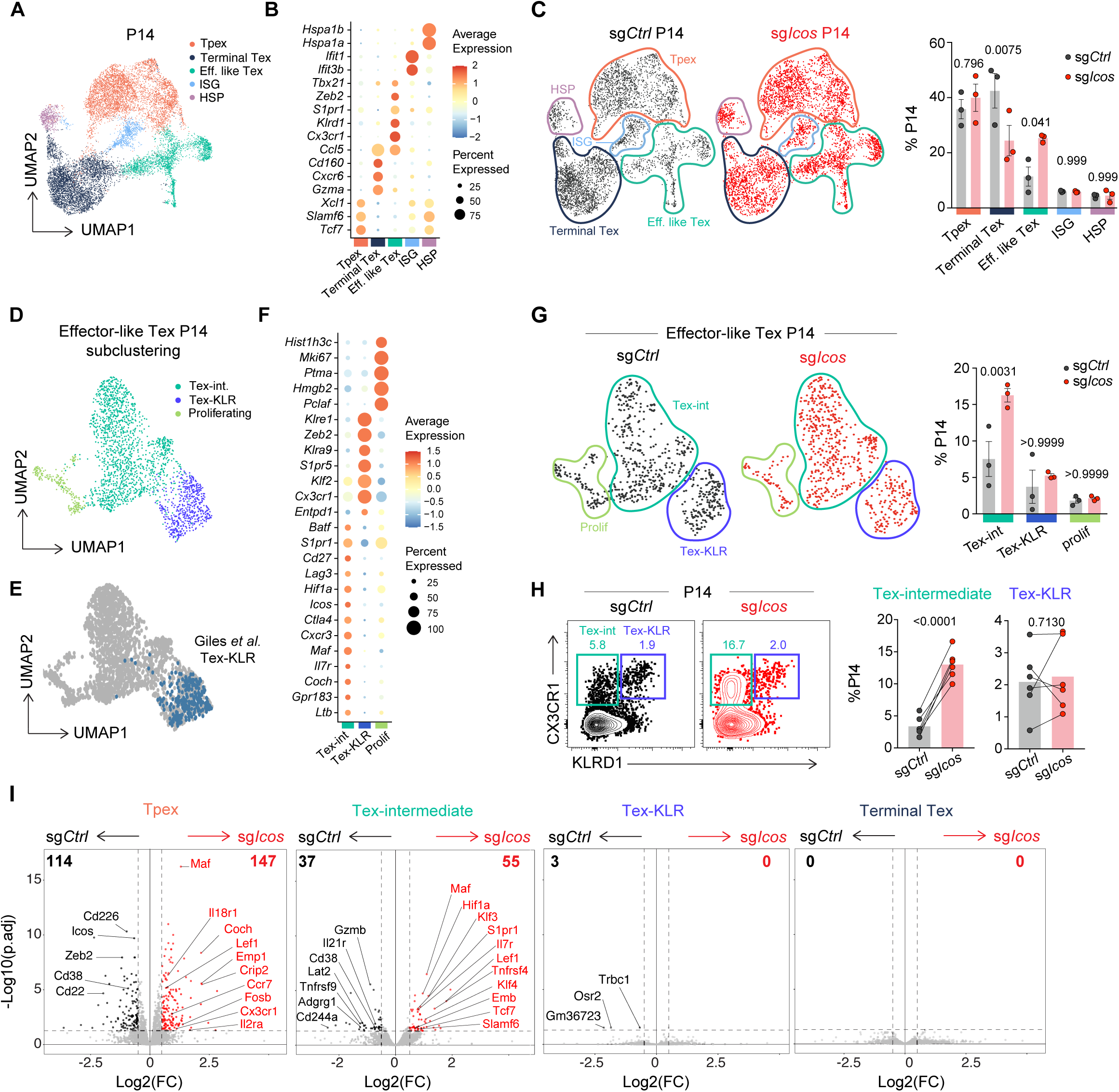
ICOS deficiency promotes accumulation of effector-like Tex. sgCtrl and sgIcos P14 were transferred into recipient mice before LCMV cl13 infection. Five weeks post infection, we performed single cell RNA sequencing (scRNA-seq) on sorted P14 (**layout Fig.S3A**). **(A)** Uniform Manifold Approximation and Projection (UMAP) of scRNA-seq clusters in sgCtrl and sgIcos P14. **(B)** Dot plot representing average and percent expression of selected genes. **(C)** UMAP of control (black) and ICOS-deficient (red) P14, graph of relative distribution of clusters. **(D)** UMAP of effector-like Tex P14 subclustered. **(E)** UMAP showing enrichment of gene signature associated with Tex-KLR^44^ **(F)** Dot plot showing average and percent expression of selected genes. **(G)** UMAP of control (black) and ICOS-deficient (red) effector-like Tex P14, graph of relative distribution of clusters. **(H)** Contour dot plots and graphs show frequency of CX3CR1^+^ KLRD1^neg^ Tex-Int and CX3CR1^+^ KLRD1^+^ Tex-KLR among control (black) and ICOS-deficient (red) P14. **(I)** Volcano plot showing differentially expressed genes (DEG) between control and ICOS-deficient Tpex, Tex-Int, Tex-KLR and terminal Tex pseudobulked profiles. Genes downregulated in ICOS-deficient P14 are shown in black, genes upregulated are shown in red. Data in (A-I) are from one experiment with three mice per group. Data in (H) are representative from two independent experiments with five mice per group. Symbols represent individual mice with bar showing the mean value of all animals analyzed and bars indicate SEM. (C and G) Significance was determined using one-way ANOVA with Holm-Sidak’s correction for multiple comparisons. (H) paired Student’s *t-test*.

A recent study in LCMV docile chronically infected animals, identified CD62L^+^ Tpex subset with improved stemness^43^. Since ICOS-deficiency significantly increased CD62L expression on Tpex (**Fig.2E and S2C**), to determine if differences observed in ICOS-deficient Tpex (**Fig.2**) were due to a specific enrichment of stem-CD62L^+^ Tpex, we next investigated heterogeneity among Tpex by further subclustering (**Fig. S3C**). We identified three potential subclusters, one of them with higher expression of *Sell* and *Myb,* similar to the previously described stem-CD62L^+^ Tpex (**Fig.S3C**). The second subcluster was characterized by higher expression of chemokines (*Xcl1, Ccl3 and Ccl4*) and the third subcluster by expression of *Il7r* and *Il18r1* (**Fig.S3C**). However, at the protein level, CD62L and IL7R did not define distinct Tpex subpopulations, and XCL-1 production was strongly correlated with IL7R expression (**Fig.S3C**). Importantly, by scRNA-seq, we did not find differences in the distribution of control and ICOS-deficient cells among the three Tpex subclusters (**Fig.S3D**).

Next, we subclustered effector-like Tex to examine heterogeneity within this subset (**Fig.3D**). Similar to previous studies in helped LCMV chronic infection^44,45^, we identified a subcluster of Tex-KLR with high expression of NK-like markers and effector molecules, as well as an intermediate subcluster, Tex-int (**Fig.3D-F**). In addition, there was also a small subcluster of proliferating cells (**Fig.3D and F**). When we compared the frequency of control and ICOS-deficient P14 between effector-like Tex subclusters, there were no differences among Tex-KLR, but ICOS-deficient P14 were specifically enriched among Tex-Int (**Fig.3G**). By flow cytometry, we confirmed increased frequency of Tex-int CX3CR1^+^ KLRD1^neg^ among ICOS-deficient P14 compared to control (**Fig.3H**), whereas we observed comparable percentages of Tex-KLR (**Fig.3H**). We next performed pseudo-bulk RNA analysis to identify differentially expressed genes in control and ICOS-deficient Tpex, Tex-int, Tex-KLR and terminal Tex (**Fig.3I**). Consistent with ICOS expression levels, we found the highest number of genes affected by loss of ICOS in Tpex (**Fig.3I and S3E**). ICOS-deficiency also altered gene expression in Tex-Int, however, very minimal changes were observed in Tex-KLR and terminal Tex (**Fig.3I and S3E**). Based on adoptive transfer experiments, Tex-int were previously reported as a transitory population capable of differentiation into Tex-KLR or terminal Tex^44,45^. Our data shpw that ICOS-deficiency did not impact Tex-KLR (numbers or phenotype), suggesting that ICOS signaling may not modulate Tex-KLR differentiation. However, we observed lower representation of ICOS-deficient P14 among the terminal Tex cluster (**Fig.3C**), indicating that in the absence of ICOS, accumulation of Tex-int might reduce or delay conversion into terminal Tex. Together, our scRNA-seq analysis highlights a prominent role for ICOS in transcriptional regulation of Tpex, as well as effector-like intermediate Tex.

### ICOS-deficiency improves survival of effector-like PD-1^+^ CD8 T cells

To understand the mechanism leading to accumulation of effector-like Tex in absence of ICOS signaling, we performed GSEA of control and ICOS-deficient effector-like P14. Surprisingly, we identified many pathways involved in cell cycle progression downregulated in ICOS-deficient effector-like Tex (**Fig.4A**). This was confirmed by lower frequency of ICOS-deficient effector-like P14 cells with recent history of proliferation, assessed by Ki67 expression (**Fig.4B**). Since accumulation of effector-like cells did not seem to result from increased proliferation, we next hypothesized that ICOS-deficient cells may have greater survival capacity. GSEA showed an enrichment of genes in the apoptosis pathway in effector-like control cells compared to ICOS-deficient (**Fig.4C**). Additionally, ICOS-deficient effector-like P14 expressed higher mRNA levels of the anti-apoptotic molecule Bcl2 (**Fig.S4A**). At the protein level, control Tpex and terminal Tex displayed higher Bcl2 expression than effector-like P14 (**Fig.4D-E**). However, ICOS-deficient effector-like cells preserved high Bcl2 levels, and overall Bcl2 expression was higher in all ICOS-deficient P14 subsets (**Fig.4D**). To directly test the sensitivity of PD-1^+^ CD8 T cells to apoptosis we cultured splenocytes from chronically infected animals for 2h *ex vivo*. We detected a higher frequency of apoptotic cells by active Caspase 3 and 7 (Cas3/7 FLICA^+^) in control P14 compared to sgIcos P14 (**Fig.4E**). Similarly, the frequency of live cells was significantly higher for ICOS-deficient P14 after three days of *ex vivo* culture without stimulation (**Fig. S4B**).

**Fig. 4:**
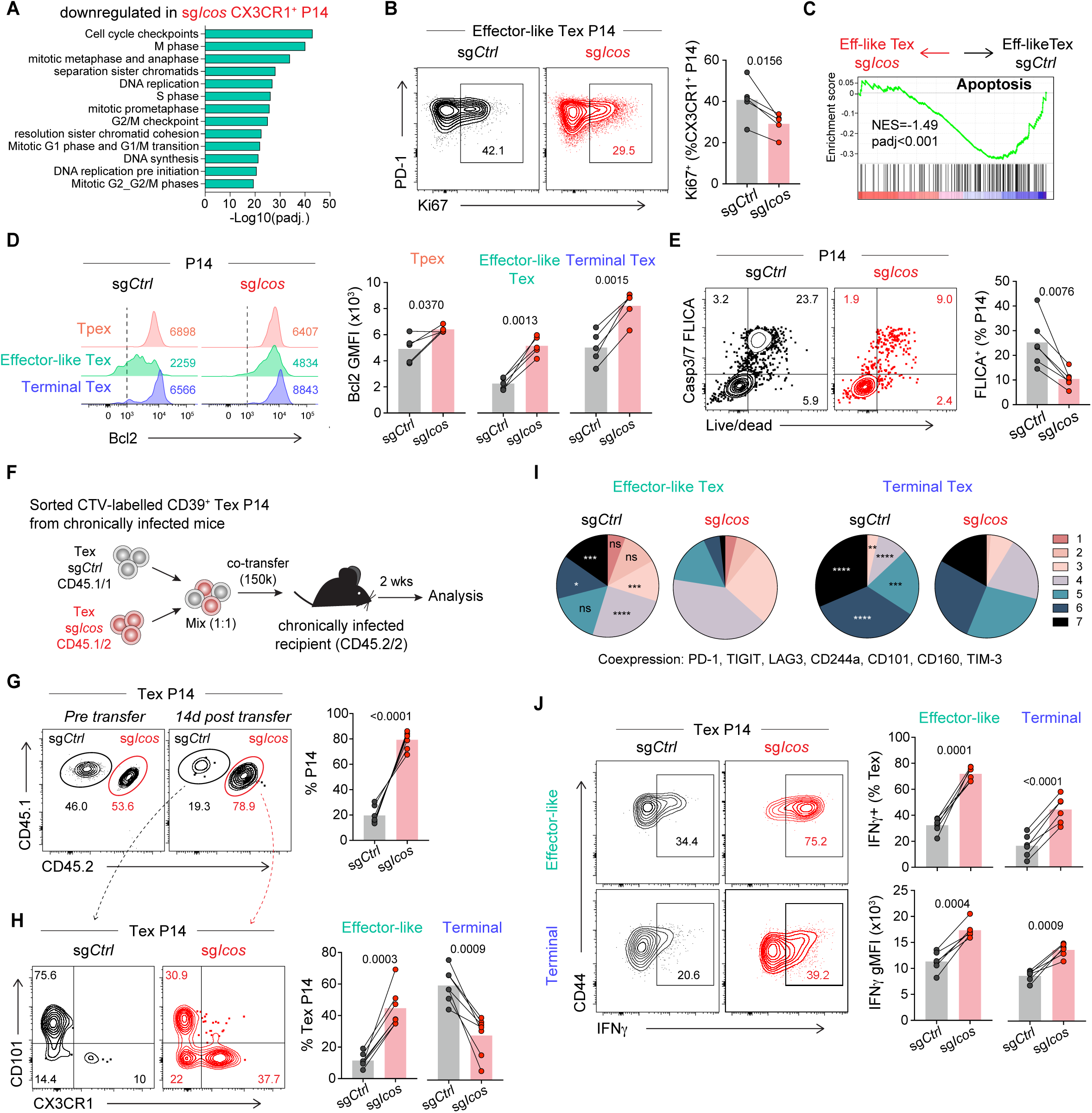
ICOS deficiency improves Tex survival and functionality. **(A)** sgCtrl and sgIcos Effector-like Tex (CX3CR1^+^ CD39^+^) P14 were sorted from spleens of LCMV-chronically infected mice and RNA was isolated for transcriptional analysis. GSEA shows top 10 significantly (adjusted P-value) downregulated Reactome pathways in ICOS-deficient CX3CR1^+^ P14. **(B)** Proliferation history (Ki67^+^) of splenic co-transferred effector-like Tex P14 sgCtrl (black) and sgIcos (red), 44 days post LCMV cl13 infection (DPI). **(C)** GSEA shows Reactome apoptosis gene set in sgCtrl versus sgIcos effector-like Tex P14. **(D)** Representative histograms and graphs show expression of anti-apoptotic protein Bcl2 in splenic co-transferred sgCtrl and sgIcos progenitor, effector-like, and terminal Tex P14 (44 DPI). Vertical dashed line indicate positivity. (**E)** Contour plots and graph show caspase 3 and 7 activities with Fluorochrome Inhibitor of Caspases (FLICA). **(F)** Experimental layout for G-H: CD39^+^ sgCtrl and sgIcos Tex P14 were sorted from chronically infected animals (28DPI), CTV-labeled, mixed 1:1 and co-transferred back into chronically infected B6 recipients (>100DPI). **(G)** Contour plots and graph show relative P14 frequency in spleen of recipient mice. **(H)** Contour plot and graphs show frequencies of effector-like Tex (CX3CR1^+^) and terminal Tex (CD101^+^) among P14 in recipient animals. **(I)** Number of inhibitory receptors (PD-1, TIGIT, LAG3, CD244a, CD101, CD160, TIM-3) co-expressed by splenic effector-like or terminal Tex sgCont and sgIcos P14 (60 DPI). (**J**) Contour plots and graphs show frequency and gMFI of IFNγ producing cells among splenic effector-like and terminal Tex sgCtrl and sgIcos P14 (60 DPI) after stimulation with GP33 peptide. Data in (B-E and I-J) are representative from 3 independent experiments with five to six mice per group. Data in (G-H) are representative of 2 independent experiments with four to seven mice per group. Symbols represent individual mice with bar showing the mean value of all animals analyzed. Connecting lines between symbols link cells analyzed from the same mouse, with bar showing the mean value. (B,D,E,G, H,J) paired Student’s t-test; (I) 2-way ANOVA with Sidak’s correction for multiple comparisons.

To directly address the persistence and survival of Tex in vivo, we sorted CD39^+^ P14 control and ICOS-deficient from chronically infected animals, then mixed them at a 1:1 ratio before transfer into chronically infected congenically distinct recipients (**Fig.4F**). Two weeks post transfer, we observed that ICOS-deficient Tex represented 80% of the transferred P14 (**Fig.4G**). Furthermore, after transfer, ICOS-deficient Tex maintained a high frequency of effector-like Tex (**Fig.4H**). Even when we accounted for differences in subset distribution pre-transfer, our data show higher persistence of ICOS-deficient effector-like cells (**Fig.S4C**).

Effector-like Tex express low levels of ICOS (**Fig.S1F**). To address if improved survival capacity is due absence of ICOS signaling in Tex or because effector-like cells emerged from ICOS-deficient Tpex with enhanced memory-like features, we performed another Tex transfer experiment, but recipient animals were treated with αICOSL antibody (**Fig.S4D**). If ICOS costimulation in Tex was responsible for lower survival of effector-like cells, preventing ICOS signaling should enhance survival of control Tex. However, two weeks post-transfer we observed a similar outcome as previous experiments, with sgIcos Tex representing 90% of the transferred cells (**Fig.S4F**). These data show that intrinsic ICOS signaling in Tex does not impact survival. Instead, it demonstrates that ICOS-deficient Tpex give rise to effector-like Tex with enhanced survival capacity, resulting in accumulation of ICOS-deficient effector-like Tex.

### Improved functionality of ICOS-deficient Tex

To evaluate potential differences in functionality of Tex in the absence of ICOS signaling, next we analyzed expression of inhibitory receptors. We observed that ICOS-deficient CD39^+^ Tex had reduced expression of checkpoint inhibitors 2B4/CD244a, CD160, and CD101 (**Fig.S4F,G**), resulting in significantly lower coexpression of inhibitory receptors by both ICOS-deficient effector-like and terminal Tex (**Fig.4I**). Furthermore, ICOS-deficient Tex had higher expression of positive costimulatory molecules, such as CD28 and OX40, which are important for T cell functionality, expansion, and metabolism (**Fig.S4H**). In addition to differences in positive and negative costimulatory receptors, we observed improved IFNγ and TNFα production by ICOS-deficient Tex (**Fig.4J and S4I**). IFNγ was increased in ICOS-deficient Tex, both in the frequency of producing cells and levels - higher IFNγ production per cell assessed by geometric Mean Fluorescence Intensity among IFNγ^+^ cells (**Fig.4J**). The combination of improved survival and enhanced capacity to produce IFNγ resulted in accumulation of nearly 10 times more IFNγ^+^ P14 when we compared ICOS-deficient cells to control in the same mice (**Fig.S4J**). Therefore, limiting ICOS signaling results in a substantial increase in the functionality of virus-specific CD8 T cells during chronic infection.

### ICOSL blockade improves Tpex memory-like phenotype and Tex functionality

ICOS is rapidly upregulated in T cells following antigen stimulation^46^, thus we cannot exclude early differences in priming accounting for the effects observed in ICOS-deficient P14. To address the role of the ICOS/ICOSL pathway on PD-1^+^ CD8 T cells without interfering with initial T cell activation, we treated mice with established LCMV chronic infection with ICOSL blocking antibodies or Ig control (**Fig.5A**). We observed increased frequency and number of virus-specific CD8 T cells in blood, spleen, lung, and liver of mice treated with anti-ICOSL (**Fig.5B and S5A-C**). Similar to genetic deficiency, ICOSL blockade improved accumulation of effector-like Tex (**Fig.5C-E and S5D**). Preventing ICOS signaling, after CD8 T cell priming, also resulted in improved memory-like phenotype by Tpex, with increased frequency of CD62L^+^ or IL7R^+^ Tpex, as well as a higher expression of LEF1 and Bcl2 (**Fig.5F**). In contrast, (but in accordance with our previous results), blocking ICOS signaling decreased expression of Tpex Tfh-associated marker Bcl6 and PD-1 (**Fig.S5E**). Finally, preventing ICOS signaling also improved Tpex cytokine production (**Fig.5G**). Together, these data indicate that ICOS can be targeted to modulate Tpex memory-like phenotype and function.

**Fig. 5:**
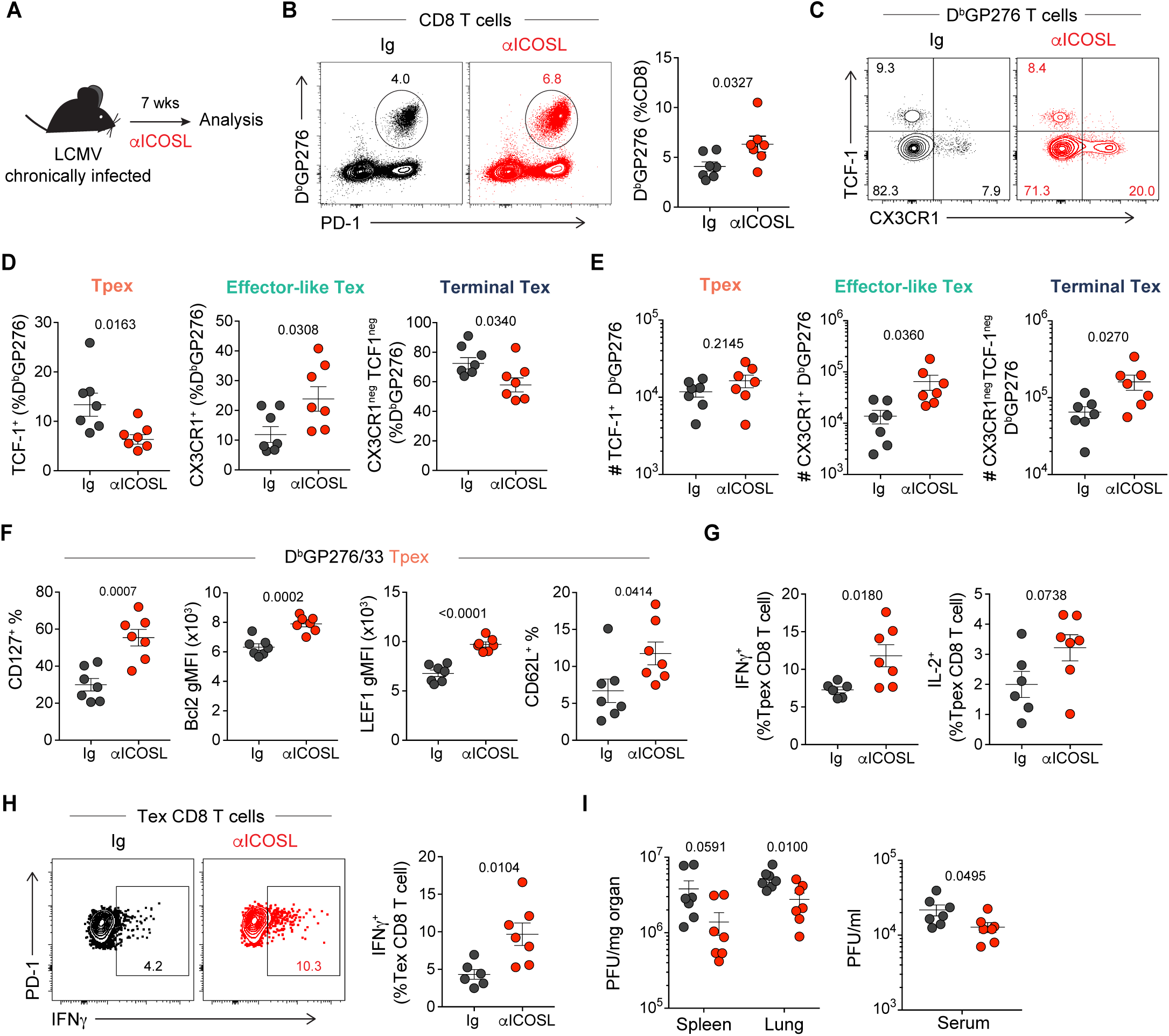
ICOSL blockade improves Tpex memory-like phenotype and Tex functionality. **(A)** Experimental layout: Mice with established LCMV chronic infection were treated with control Ig or ICOSL blocking antibodies (αICOSL) for 7 weeks. B-H show analysis in the spleen after treatment. (B) Contour plots and graph show frequency of LCMV-specific CD8 T cells. **(C-D)** Frequency and **(E)** absolute number of TCF1^+^ progenitor, CX3CR1^+^ effector-like, and CX3CR1^neg^TCF-1^neg^ terminal Tex LCMV-specific (D^b^GP276) CD8 T cells. **(F)** Graphs show frequency or gMFI of memory markers expressed by Tpex LCMV-specific CD8 T cells. **(G)** Frequency of IFNγ and IL-2-producing TCF-1^+^ PD-1^+^ CD8 T cells after LCMV peptides restimulation. **(H)** Contour plots and graph show frequency of IFNγ-producing CD39^+^ PD-1^+^ CD8 T cells after LCMV peptides restimulation. **(I)** Viral load in spleen, lung, and serum 7 weeks post-treatment. Data in (B-I) are representative from 2 independent experiments with five to seven mice per group. Symbols represent individual mice. Data are shown as mean +/- SEM. (B-I) unpaired Student’s t-test.

Next, we further analyzed the effects of ICOSL blockade in Tex. Preventing ICOS signaling improved Bcl2 expression in effector-like virus-specific CD8 T cells **(Fig.S5F**), while also lowering expression of inhibitory receptors CD101, CD244a and CD160 (**Fig.S5G**). Blocking ICOSL not only increased effector-like virus-specific CD8 T cells, but also improved IFNγ production by virus-specific Tex (**Fig.5H**, **FigS5H**). Overall, preventing ICOS signaling enhanced accumulation of IFNγ-producing virus-specific T cells (**Fig.S5I**) and resulted in better viral control in lung and serum of chronically infected animals (**Fig.5I**).

### ICOS regulates functionality of tumor-specific CD8 T cells

To address to what extent ICOS can modulate tumor-specific CD8 T cell differentiation and function, we used a modified genetically engineered mouse model of liver cancer that resembles hepatocellular carcinoma (HCC)^47^. Hydrodynamic injection of oncogenic plasmids induces hepatocyte transformation, through p53 deletion and overexpression of Myc, as well as expression of luciferase (Luc) to monitor tumor growth and expression of the model antigen Ovalbumin (OVA)^47^ (**Fig.S6A**). The original construct (LucOVA-SIY) contained two major CD8 T cell epitopes, SIYRYYGL (SIY) and SIINFEKL (OT-I epitope, OVA257–264)^47^. To decrease immunogenicity and improve tumor penetrance, we removed the SIY epitope and mutated SIINFEKL into SI**Y**NFEKL (Y3) to reduce OT-I sensitivity for OVA peptide^48^. We transferred a small number of tumor-specific CD8 T cells (OT-I; OVA-specific CD8 T cells) into mice 3 days after receiving oncogenic plasmids encoding Luc-OVAY3 **Fig.6A**). Four weeks post hydrodynamic injection, OTI in liver differentiated into Tpex (PD-1^+^ TCF-1^+^ SLAMF6^hi^) as well as CD39^+^ CXCR6^+^ Tex (**Fig.6B**). Similar to LCMV chronic infection, the majority of Tpex expressed ICOS while only 20% of differentiated Tex were ICOS^+^ (**Fig.6C**). Tumor burden is a major driver of CD8 T cell differentiation; to address the intrinsic role of ICOS, we next co-transferred sg*Ctrl* and sg*Icos* OT-I into mice that had received hydrodynamic injection with oncogenic plasmids to phenotype tumor-specific CD8 T cells in a similar tumor microenvironment (**Fig.6D**). Three- and four-weeks post-tumor initiation, we observed similar frequency of control and ICOS-deficient OT-I in tumors (**Fig. S6B**). Interestingly, the relative frequency of sg*Icos* Tpex was preserved with tumor progression contrary to sgCtrl OT-I (**Fig.6E**), suggesting improved maintenance of Tpex in the absence of ICOS signaling. Additionally, sg*Icos* Tpex displayed increased CD127 expression (**Fig.6F**), and improved capacity to produce IFNγ and IL-2 (**Fig.S6C**). We also observed increased IFNγ production by ICOS-deficient Tex compared to control OTI (**Fig.6G**). Differences in cytokine production were more pronounced at d30, suggesting improved capacity of ICOS-deficient cells to preserve functionality over time.

**Fig 6.**
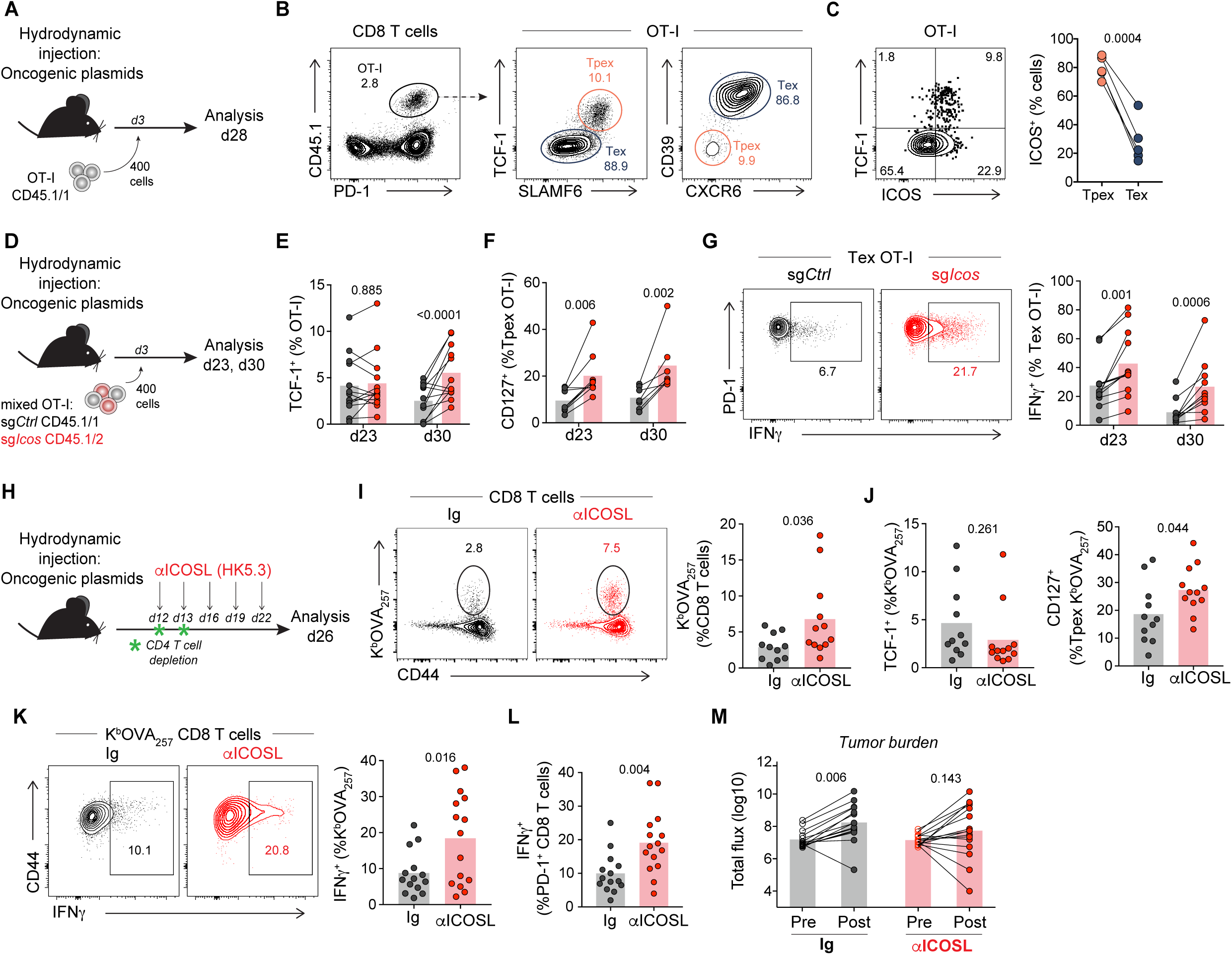
ICOS regulates the functionality of tumor-specific CD8 T cells. **(A)** Experimental layout for (B-C): Mice received hydrodynamic tail vein injection (HDTVI) with oncogenic plasmids and 3d later, 400 OT-I were transferred. Livers were harvested and analyses were performed 28d post tumor initiation. **(B)** Representative plots and gating strategy of TCF-1^+^ SLAMF6^+^ Tpex and CD39^+^ CXCR6^+^ Tex OT-I infiltrating HCC. **(C)** ICOS expression on Tpex and Tex OT-I. **(D)** Experimental layout for (E-G): 400 total sgCtrl and sgIcos transfected OT-I were cotransferred 1:1, 3d post HDTVI. Analyses were performed three and four weeks post tumor initiation. **(E)** Frequency of Tpex (TCF-1^+^) OT-I. **(F)** CD127 expression on Tpex OT-I. **(G)** Representative plot (d23) and frequency of IFNγ producing Tex OT-I after 3h of PMA/ionomycin stimulation. **(H)** Experimental layout for (I-M): Mice were depleted of CD4 T cells prior to treatment with Ig control or ICOSL blocking antibodies starting 12d post HDTVI of oncogenic plasmids. Ig or αICOSL was administered every 3d for 2 weeks before analysis. **(I)** Frequency of K^b^OVA_257_ tumor-specific CD8 T cells. **(J)** Frequency of Tpex K^b^OVA_257_ and CD127 expression by K^b^OVA_257_ Tpex. **(K-L)** IFNγ production by **(K)** K^b^OVA_257_ cells and **(L)** PD-1^+^ CD8 T cells after PMA/ionomycin stimulation. **(M)** Total flux (photons/sec) as measured by bioluminescence imaging pre and post treatment. Data in (A-C) are representative of 3 independent experiments, (D-M) combined data from 2 independent experiments with five to eight mice per group. Symbols represent individual mice. Connecting lines between symbols link cells analyzed from the same mouse. Data are shown as mean. (E-G) paired Student’s t-test; (I-L) unpaired Student’s t-test; (M) ANOVA with Sidak’s correction for multiple comparisons.

We next sought to address how ICOSL blockade would impact tumor control. Twelve days post hydrodynamic injection, mice were stratified based on luciferase detection by In Vivo Imaging System (IVIS) into groups either receiving Ig control or ICOSL blockade for two weeks (**Fig.6H**). Since ICOS signaling is important for both regulatory and conventional CD4 T cell function, mice were depleted of CD4 T cells, in order to specifically assess the impact of ICOS signaling in CD8 T cells (**Fig.6H**). ICOSL blockade led to increased frequency of tumor-specific CD8 T cells in tumors (**Fig.6I** and **Fig.S6D**). ICOSL blockade also enhanced Tpex CD127 expression (**Fig.6J**) as well as their cytokine production (**Fig.S6E**). Next, we confirmed that ICOSL blockade improved IFNγ production by tumor-specific CD8 T cells (**Fig.6K-L**). Notably, ICOSL blockade resulted in a slower tumor growth in some of the animals (**Fig.6M**). Overall, our data suggest that blocking ICOS signaling on tumor-specific CD8 T cells improves functionality and tumor control.

## DISCUSSION

In this study, we address the function of ICOS in the differentiation of exhausted PD-1^+^ CD8 T cells. During antigen persistence, Tpex maintain antigen-specific CD8 T cell responses. Tpex differentiation into Tex, which retain residual effector function, is essential for virus or tumor control. However, the mechanisms leading to efficient Tpex differentiation and maintenance of effector function in Tex remain unclear. Tpex express high levels of costimulatory molecules, that can modulate their differentiation and survival^18^. Here we uncovered that ICOS signaling in PD-1^+^ CD8 T cells restrains accumulation of effector-like Tex. Our data show that lack of ICOS signaling increases FoxO1 activity and improves memory gene expression programs by Tpex. We find that ICOS-deficient Tpex can give rise to effector-like Tex with improved survival capacity and functionality. Furthermore, we demonstrate that targeting ICOSL with blocking antibodies can improve functionality of PD-1^+^ CD8 T cells, leading to better viral control in chronic infection and delay of tumor growth in a mouse model of HCC. Overall, our data show that sustained ICOS costimulation is detrimental for the efficacy of antigen-specific CD8 T cells during antigen persistence.

Mechanistically we show that ICOS signaling modulates FoxO1 localization in Tpex. Previous studies had demonstrated that FoxO1 activity is critical for the acquisition of stem-like properties by T cells^49^ and required for naïve T cell homeostasis^50^ as well as development and maintenance of memory T cells^27–30^. During antigen persistence, FoxO1 was shown to be required for Tpex survival and development of exhausted CD8 T cell population^31,32^. Here we find increased FoxO1 nuclear retention and transcriptional changes consistent with enhanced FoxO1 activity in ICOS-deficient Tpex. Loss of ICOS by Tpex promoted memory-associated gene expression programs and improved functionality of exhausted T cells. Echoing our results, recent reports demonstrated that FoxO1 overexpression in chimeric antigen receptor (CAR) T cells was associated with increased memory features leading to improved persistence, functionality and therapeutic efficacy^36,37^. Similarly, our results suggest that, improving FoxO1 activity in chronically stimulated CD8 T cells reinforces progenitor stemness and improves functionality. Of note, Doan *et al.*, show that restricting FoxO1 to the nucleus (by mutations that prevent phosphorylation) is detrimental for CAR T cells, due to inhibition of differentiation, hampering acquisition of effector function. Overall these data suggest that enhancing FoxO1 activity is beneficial, but it needs to be regulated to allow for T cell differentiation. Our data suggest that optimal FoxO1 activity may be achieved by limiting ICOS signaling in PD-1^+^ CD8 T cells.

Effector-like Tex are a transitory population that eventually differentiates to a more dysfunctional terminal Tex stage^7,44,45^. Differentiation of Tpex into effector-like Tex is essential for viral or tumor control^7,8^. CD4-help and PD-1 targeted therapy increase effector-like Tex^7,8^. Efficacy of immunotherapy relies not only on increased number of antigen-specific CD8 T cells but also on accumulation of CD8 T cells with improved functionality. In the absence of ICOS signaling, we observed more effector-like Tex, even in the absence of CD4 T cell help. We demonstrated that transitory cells derived from ICOS-deficient Tpex had improved survival capacity and functionality, and delayed progression toward terminal Tex. Furthermore, ICOSL blockade promoted accumulation of effector-like Tex with enhanced functionality, resulting in improved viral control. Similarly, ICOSL blockade in HCC, improved cytokine production by tumor-specific Tex and delayed tumor growth. Thus, we propose a new approach to reinvigorate antigen-specific CD8 T cell responses, by targeting and improving stemness of progenitor exhausted CD8 T cells leading to enhanced and more sustainable effector-like responses.

Recent work by Peng et al. highlighted the requirement of ICOS signaling for the development and maintenance of tissue resident memory (Trm) CD8 T cells after acute infection^22^. Mechanistically, ICOS-deficient CD8 T cells failed to efficiently downregulate the transcription factor KLF2, responsible for expression of the trafficking receptor S1PR1. SIPR1 downregulation is required to establish Trm^51^. In LCMV chronic infection, Tpex reside in lymphoid organs^52^, whereas in tumors, Tpex can be found within APC-rich niches^11,12^. Importantly, we did not observe defects in the residency of ICOS-deficient virus-specific PD-1^+^ CD8 T cells during chronic infection. In fact, we observed enhanced localization of ICOS-deficient virus-specific CD8 T cells in the white pulp of the spleen. However, in line with previous studies, ICOS deficiency improved *S1pr1* expression and we observed an increased frequency of circulating effector-like Tex.

ICOS-L is expressed by antigen presenting cells^23^, but also non-hematopoietic cells and tumor cells^19–22^. Interestingly, a bystander contribution of ICOS engagement was described for the maintenance of follicular helper CD4 T cells^53^ and resident memory CD8 T cells^22^. Future work will need to determine the contribution of bystander ICOS signaling for chronically activated PD-1^+^ CD8 T cells, especially in inflammatory contexts, since ICOS-L can be upregulated in non-hematopoietic cells by TNFα^19^.

ICOS is rapidly upregulated following naïve T cell activation, and the majority of effector CD8 T cells express high levels of ICOS^54^. Our data revealing the negative role of ICOS on PD-1^+^ CD8 T cells responses is paradoxical given the general positive function of costimulation, including our previous data on CD28^18^, and even previous studies on ICOS. For example, in a model of adoptive cell therapy ICOS-deficient tumor-specific CD8 T cells showed lower cytotoxic capacity and tumor control^54^. Furthermore, in acute listeria infection, ICOS overexpression in CD8 T cells leads to increased cytolytic activity^55^. However, ICOS overexpression skews CD8 T cell differentiation toward short-lived effector cells (SLEC) at the expense of memory precursor, resulting in lower capacity to produce cytokine and mount protective memory response^55^. These data suggest a role for ICOS into SLEC terminal differentiation rather than polyfunctional memory cells. Our data show that ICOS signaling is detrimental for acquisition of memory properties by Tpex and subsequently limits fitness of differentiated PD-1^+^ CD8 T cells, since loss of ICOS improves survival and ability to produce cytokines by effector-like Tex. Hence, our study highlights an unappreciated detrimental role of sustained ICOS costimulation for PD1^+^ CD8 T cells during chronic antigen exposure.

The widespread expression pattern of ICOS may also account for some contradictory reports regarding benefit of ICOS engagement. It is critical to consider that ICOS is highly expressed by regulatory T cells and supports their immunosuppressive function through IL-10 production^21,56,57^. Previous work had shown that ectopic expression of ICOSL by tumor cells enhanced immunosurveillance and limited tumor growth^58–61^. In addition, work by Fan *et al.* have demonstrated a synergy between agonistic ICOS costimulation and Cytotoxic T-lymphocyte associated protein 4 (CTLA-4) blockade in subcutaneous tumor models^58^. In this preclinical model, enhancement of ICOS signaling during anti-CTLA4 therapy improved Th1 effector responses as well as M1 macrophage polarization^58,62^. Furthermore, ICOS signaling is essential for Tfh differentiation and function^24–26^. We and other have shown that presence of CD4 T cells with Tfh features in solid tumors is associated with positive clinical outcomes^12,63,64^. Yet, our results show that ICOS signaling is detrimental for PD-1^+^ CD8 T cells during persistent antigen exposure. Importantly, we address ICOS functionality in a CD8 T cell-intrinsic manner. For example, we depleted CD4 T cells during ICOSL blockade in HCC bearing mice to study effects in CD8 T cells. Reconciling our findings with previous studies, work by Yu et al. in graft-versus-host disease (GVHD), highlighted opposing effects of ICOS on CD4 and CD8 T cells. Whereas ICOS-deficient CD4 T cells demonstrated impaired effector function and lower induction of GVHD pathology, ICOS-deficient CD8 T cells significantly worsen GVHD morbidity^65^. Similar to our results, there was improved survival capacity and cytokine production by ICOS-deficient CD8 T cells^65^. Our findings add another layer of complexity that may prevent success of broad ICOS agonist therapy, with seemingly opposite requirements for ICOS costimulation on CD4 and CD8 T cell function during chronic antigen exposure.

Preclinical studies have highlighted the potential of agonistic ICOS in combination with PD-1 targeted therapy to improve cancer immunotherapy^66^. However, recent updates report modest clinical activity of agonistic ICOS in combination with nivolumab (anti-PD-1) in cancer patients^67,68^. Differential cell-specific requirement for ICOS costimulation may account in part for these observations. It is also critical to take into consideration cellular mechanisms and targets of immune checkpoint blockade to select ideal therapies to combine with strategies targeting the ICOS/ICOSL pathway. Whereas PD-1 targeted therapy has a stronger impact on the CD8 T cell compartment, anti-CTLA-4 has a major effect on CD4 T cells^69^. Thus strategies aiming to improve ICOS signaling would be more beneficial combined with therapies involving CD4 T cell activation such as anti-CTLA4. Nevertheless, it would be important to design therapies specifically targeting ICOS on CD4 T cells. Similarly, strategies that limit ICOS costimulation on PD-1^+^ CD8 T cells, while preserving ICOS signaling on other cell types, might be valuable for therapeutic efficacy aiming to enhance CD8 T cell responses.

Overall, our work supports the concept that sustained ICOS costimulation is detrimental for PD-1^+^ CD8 T cells. Our data show that limiting ICOS signaling improves FoxO1 activity and stemness of progenitor cells, leading to enhanced functionality of PD-1^+^ CD8 T cells. We report similar results in chronic viral infection and a mouse model of HCC, showing that ICOSL blockade leads to better viral or tumor control. Here we demonstrate that promoting memory-like properties at the expense of Tfh transcriptional program is beneficial for Tpex, with major implications for therapies aiming to improve PD-1^+^ CD8 T cell responses. Our data suggest a new approach to reinvigorate PD-1^+^ CD8 T cells, by targeting and improving stemness of progenitor cells to enhance effector-like responses and functionality of exhausted PD-1^+^ CD8 T cells.

### Limitations of the study

While our data show that ICOS deficiency increases FoxO1 activity in Tpex, there are additional transcriptional differences between control and ICOS-deficient Tpex that may contribute to the observed phenotype. However, identifying ICOS-specific effects that are FoxO1-independent becomes challenging since FoxO1 is essential for survival of PD1^+^ CD8 T cells^31,32^. Moreover, forcing FoxO1 nuclear localization to increase its activity is not a suitable approach either, as FoxO1 regulation must remain dynamic to allow for its nuclear exclusion and Tex differentiation^36^. Finally, even though we did not observe that ICOS deficiency reduced initial priming/expansion of CD8 T cells, it is conceivable, that in situations of sub-optimal priming or low antigen abundance, ICOS may enhance initial activation of CD8 T cells.

## Acknowledgments

We thank the NIH Tetramer Core Facility, Center for Comparative Medicine and Surgery, BioMedical Engineering and Imaging Institute, Dean’s Flow Cytometry CORE and Center for Advanced Genomics Technology at the Icahn School of Medicine at Mount Sinai (ISMMS). Oncological Sciences Sequencing Core at the ISMMS. We thank Arun Narasimhan from the Microscopy CORE at ISMMS for his help with immunofluorescence image analysis. We thank Travis Dawson, Darwin D’souza and Krista Angeliadis from Human Immune Monitoring Center at the ISMMS for their help with single cell RNA sequencing. We thank Pauline Hamon for sharing the hepatocellular carcinoma digestion protocol. A. Lujambio was supported by Damon Runyon-Rachleff Innovation Award (DR52-18) and R37 Merit Award (R37CA230636). Funding was provided by the NIH (R01 AI153363-01A1 to A.O.K., E.H., I.K., J.L.) and Merck & Co Inc (MK3475, IISP 61083 to A.O.K., B.R.R., A.V.).

## Author contributions

Conceptualization, E.H., I.K., A.O.K.; Methodology, E.H., I.K., A.V., A.O.K.; Formal Analysis, E.H., I.K., N.P., D.F., T.B., B.R.R; Investigation, E.H., I.K., J.L., A.V., N.P., V.v.d.H., M.D.P.; Validation, T.B., B.R.R.; Resources A.M., E.B., B.D.B., D.D.S.; Writing - Original draft, E.H., I.K., A.O.K.; Writing – Review & Editing, all authors; Visualization, E.H., I.K.; Supervision, E.H., A.O.K., D.D.S., B.R.R.; Project Administration, E.H., A.O.K.; Funding Acquisition, A.O.K.

## Data and materials availability

Raw data from bulk RNA-seq experiment have been deposited at GEO, accession number GSE270420. Raw and demultiplexed data from scRNA-seq experiment have been deposit at GEO, accession number GSE270606. All reagents generated in this study are available from the Lead Contact. All data needed to evaluate the conclusions in the paper are present in the paper.

## METHODS

### Mice, infections and hydrodynamic injections

C57BL/6 mice were purchased from the Jackson Laboratory (C57BL/6J, stock 000664) or Envigo (C57BL/6NHsd). Congenic B6 CD45.1 (B6.SJL-*Ptprc^a^Pepc^b^*/BoyJ, stock 002014) mice were purchased from the Jackson Laboratory. CD45.1 P14 TCR transgenic mice^70^ (TCR specific for the H2-D^b^-restricted LCMV glycoprotein epitope GP33) were bred in house. OTI OVA TCR transgenic (stock 003831, TCR specific for the H2-K^b^-OVA_257-264_) were purchased from Jackson Laboratory and bred to B6 CD45.1 in house. Mice were infected at 6-8 weeks of age. For acute LCMV infection and development of memory CD8 T cells, mice were given 2x10^5^ plaque forming units (PFU) LCMV Armstrong (Arm) intraperitoneally (i.p.). For life-long chronic LCMV infection, the helpless model was used, unless otherwise indicated^71^: mice were given 200 μg of the CD4-depleting antibody GK1.5 i.p. (BioXcell) 1 day prior to infection and again on the day of infection with 2x10^6^ PFU LCMV clone 13 by intravenous route (i.v.). For helped LCMV infection, mice were infected with 2x10^6^ PFU LCMV clone 13 i.v.. Mice received further treatments once chronic infection was established (at least 40 days post-infection), unless otherwise noted. Viral load was assessed by plaque assays on Vero E6 cells as previously described^72^.

For HCC development, mice received hydrodynamic injection with oncogenic plasmids, as described, but pT3-EF1a-MYC-IRES-lucOY3 was generated from pT3-EF1a-MYC-IRES-lucOS^47^, gifted by Dr. Amaia Lujambio. Mice were injected at 7-8 weeks of age or at least 18g of weight. A sterile saline solution/plasmid mix was prepared containing: 15μg of pT3-EF1a-MYC-IRES-lucoY3 (Luco-Y3), 20μg of px330-sg-p53 (sgp53), and 3.75µg of SB13 transposase-encoding plasmid dissolved in 2 ml of saline solution. Mice received intravenous injection of saline/plasmid mix using 26G needle, volume corresponding to 10% of the weight of each mouse. Vectors for hydrodynamic delivery were produced by Azenta Life Sciences/Genewiz.

All animal experiments performed in this study were approved by the Institutional Animal Care and Use Committee of Icahn School of Medicine at Mount Sinai (protocol number IACUC-2018-0018/PROTO201900610).

### Naïve CD8 T cells electroporation and adoptive T cell transfer

sgRNA targeting murine *Icos* (sgIcos#1:TGGTCTTGGTGAGTTCGCAG; sgIcos#2: GCAGAAGTAATAGCTTCCCT) and *F8* (used as control gene not expressed in T cells) (sgF8#1: TCTCCAATGAAGCTTGACGG; sgF8#2: TAATAGCGGGTCAGGCACCG) were purchased from Synthego (CRISPRevolution sgRNA EZ Kit, Synthego).

For sgRNA/Cas9 RNP formation, 1 µl of sgRNA (0.3 nmol/µl in nuclease-free H2O) was incubated with 0.6µl of Cas9 (10 mg/ml, Alt-R® S.p. Cas9 Nuclease V3, Integrated DNA Technologies) in nuclease-free H2O in a final volume of 5 µl for 10 min at room temperature. Efficiency of deletion was assessed separately for both sgRNA targeting Icos and then used in combination for experiments (0.5µl of each gRNA was incubated with Cas9 for 10min).

CD8 T cells were enriched from splenocytes from naïve P14 or OT-I mice with the EasySep Mouse CD8 T Cell Isolation Kit (StemCell). 2x10^3^ P14 or OT-I were washed in PBS, resuspended in 20µl P3 buffer (P3 primary cell 4D-Nucleofector™ X Kit S, Lonza Ltd), and mixed with the 5 µl sgRNA/Cas9 ribonucleotide complex prior to electroporation using the P3 primary cell 4D-Nucleofector™ X Kit S electroporation wells and Lonza 4D-Nucleofector™ System (Pulse CM137). Immediately after electroporation 150µl of media (RPMI 10% FBS, 2mM L-glutamine, 10mM HEPES, 55µM β-mercaptoethanol) was added to each well and cells were rested for 30min at 37°C before further processing^33,34^. 1-3x10^3^ P14 CD8 T cells (or 1x10^3^ each P14 for co-transfer) were transferred i.v. into B6 recipient mice the same day of infection with LCMV clone 13. Only 400 OT-I (200 each OT-I for cotransfer) were transferred 3 days post hydrodynamic injection with oncogenic plasmid.

For P14 Tpex or Tex transfer, splenocytes were isolated from LCMV clone 13 infected mice 25-30 days post-infection. Donor mice had received 3x10^3^ P14 (control or sgIcos) before infections, as described above. After CD8 enrichment (StemCell), cells were labeled with 5 µM cell trace violet (CTV, Thermo Fisher Scientific) for 20min at 37°C. CTV staining was quenched for 5 min with RPMI containing 10% FBS before proceeding with cell surface staining with anti-CD8α (53-6.7), -CD44 (IM7), -CD45.1(A20), -CD45.2(104), -CD39 (24DMS1) and -PD-1 (RMP1-30; non-blocking clone) from BD or Biolegend and Live/Dead fixable dead cell stain (Invitrogen). Live CD8^+^ PD-1^+^ CD45.1^+^ CD45.2^+/-^ CD39^neg^ Tpex and CD39^+^ (Tex) were isolated by FACS (post-sort purity >96%). After sort, control and sgIcos P14 cells were mixed 1:1 and injected into B6 recipients (chronically infected or cleared of LCMV Arm infection, as indicated in figures). 2-8 weeks post-transfer, single cell suspensions were obtained from spleen and lung for staining and analysis.

### Single cell suspension from spleen, blood and other tissues from LCMV-infected animal

Spleens were digested for 30 minutes at 37°C with 0.4 U/ml of collagenase D (Roche, # 11088882001) in HBSS (Gibco) with 2% FBS (Gibco) then fully disrupted through 100µm strainers and subjected to red blood cells lysis (ACK, Gibco) for 3min at RT. Exception was made for CXCR3 or CXCR5 stains; spleens were not digested and were instead directly disrupted through strainer to preserve chemokine receptor expression.

Peripheral blood was collected in 400µl 4% sodium citrate tribasic. 2ml RPMI 2%FBS was added to collected blood and vortexed. Sample was underlaid with 1.5ml histopaque-1077 (Sigma) and spun at 2000rpm 20min RT without break. Interphase containing cells was collected in a new tube and washed with RPMI 2%FBS.

Before collection, lung was perfused by injection of cold PBS into the right side of the heart. Lung was cut into pieces and digested for 1h at 37°C with 150 U/ml Collagenase type I (Gibco, 17100-017). Pieces were fully disrupted through 100µm strainers and pellet resuspended in 44% Percoll (Citiva/Fisher), underlaid with 67% Percoll and spun 20min at 2000rpm, RT without brake. Interface containing lymphocytes was collected with transfer pipette into new tubes, and, after cells were pelleted, RBC lysis was performed for 3min at RT.

Liver was perfused with cold PBS injected into hepatic artery, gall bladder was removed and whole liver collected. Liver was not digested and was processed similarly to lung to obtain single cell suspension.

Axillary, brachial, inguinal and cervical lymph nodes were collected in RPMI with 10%FBS. Lymph nodes were cut into small pieces and homogenized through 100µm to obtain single cell suspension.

### *In vivo* treatments

For blockade of ICOSL molecules, 200μg anti-mouse ICOSL (HK5.3, BioXcell) were administered i.p. every 3 days. Control mice were received 200 μg isotype control Rat IgG2a, i.p. every 3 days. Final analyses were performed 2-7 weeks after treatment initiation as indicated.

### Antibodies and flow cytometry

Single cell suspensions were surface stained for 30 min on ice in flow cytometry buffer (PBS, 2% FBS, 1mM EDTA, 0.05% sodium azide) supplemented with 10% brilliant stain buffer (BD bioscience) if more than two brilliant violet dyes were used and with anti-CD16/32 (Trustain FcX, Biolegend). Biotinylated monomers Db/GP33-41, Db/GP276-286 and OVA-derived H2Kb/OVA257 were obtained from NIH tetramer core facility and tetramerized as previously described^73^. Tetramers with LCMV epitopes were stained together with surface antibodies, but K^b^OVA tetramer staining was performed before surface staining. Cells were then incubated with fixable viability dye (Thermofisher) in PBS for 10 min on ice. For intracellular cytokine staining, samples were fixed/permeabilized for 20 min at RT with the BD Cytofix/Cytoperm kit (BD Biosciences). For transcription factor staining, samples were fixed/permeabilized overnight at 4°C with Foxp3/Transcription Factor Staining Buffer Set (eBioscience). Intracellular staining was performed at RT for 45min.

For intracellular cytokine staining, 2x10^6^ splenocytes were incubated with 0.1 μg/ml of GP33 LCMV peptide for P14 stimulation or a pool of LCMV peptide (GP33, GP276, GP92, GP118, NP235, NP205), in the presence of GolgiPlug (brefeldin A) and GolgiStop (monensin) for 5h at 37°C. TILs from HCC-bearing mice were stimulated with cell activation cocktail (Biolegend, mixed phorbol 12-myristate-13-acetate and ionomycinin) in the presence of GolgiPlug (brefeldin A) and GolgiStop (monensin) for 3h at 37°C.

For *in vivo* intravascular labelling we injected intravenously 3µg of anti-CD8β.2 FITC (Biolegend). 3 to 5 min post injection, mice were euthanized and organs collected for processing and analysis

### Bulk RNA-Sequencing

10,000 cells were FACS sorted into lysis reagent (QIAGEN). After addition of chloroform (Sigma) and centrifugation, RNA was isolated from the aqueous phase using RNEasy Micro kit (QIAGEN). 1 ng RNA was used as input for the NEBNext Single Cell/Low Input RNA Library Prep Kit for Illumina (NEB). Poly-A enriched libraries were sequenced in 75bp single-end mode on a Novaseq 6000 system (Illumina) in the ISMMS NGS platform (Center for Advanced Genomics Technology). Reads were quasi-mapped to the Gencode M25 (GRCm38.p6) gene set using *salmon* 1.2.1^74^, transcripts were summarized to gene level using *tximeta* 1.14.1^75^ and differential gene expression analysis was performed between ICOS-deficient vs. control in either CD39^neg^CX3CR1^neg^ P14 Tpex or CD39^+^CX3CR1^+^ effector-like Tex or CD39^+^CX3CR1^neg^ terminal Tex P14 using *DESeq2* 1.36.0^76^. Independent hypothesis weighting (IHW) was performed on p-values with the default covariate parameter of mean counts of each gene and significance level of 0.05 for FDR control^77^. Significant genes were retained by filtering for an IHW-adjusted p-value < 0.05 and a log2 fold change ≥ 0.5 or ≤ - 0.5. GSEA in Fig.2 was performed on the entire expressed gene set with GSEA version 4.3.3 using the pre-ranked list mode, running the gene set permutation 500 times. Gene set enrichment analysis in Fig.4 was performed with the *fgsea* package^78^ on the entire expressed gene set pre-ranked based on the Wald test statistic computed by DESeq2, significant pathways were reported using an adjusted P-value cutoff of 0.05 and ordered by adjusted P-value. GSEA was also conducted on DEG using ChIP-X-Enrichment Analysis from Enrichr database (https://maayanlab.cloud/Enrichr/)^79,80^. Gene sets were used from Doan *et al.* (FoxO1 regulon)^36^, from Joshi *et al.* (memory CD8 T cells)^81^ and from Hale *et al.* (Tfh cells)^82^. For heatmaps, the Z-score was calculated as (sample TPM - mean of all TPM) divided by standard deviation.

### Confocal microscopy and analysis

FoxO1 cellular localization was performed as previously described^83^. Briefly, splenocytes from chronically infected animals transferred with P14 sgCtrl or sgIcos were surface stained with anti-CD8α (53-6.7), -CD44 (IM7), -CD45.1(A20), -CD45.2(104), -PD-1(RMP1-30) and live/dead IR, then fixed 20min at RT with IC fixation buffer (00-8222-49, eBioscience) before sort. Around 15-20k sorted Tpex (sgCtrl or sgIcos) or memory P14 (from ARM memory mouse) were further fixed for 20 minutes at room temperature with a solution containing 3.13% Glyoxal fixative (Sigma-Aldrich), 20% ethanol, 0.75% acetic acid (pH=4–5). Cytospun cells were washed in PBS, then blocked/permeabilized in PBS/0.3%BSA/0.3% Triton-X100 for 1 hour at room temperature. After overnight incubation with anti-FoxO1 antibody (C29H4, Cell Signaling Technologies, 1:200) at 4°C, slides were washed with PBS-Tween, incubated in ImmPress anti-rabbit secondary antibody for 1 hour at room temperature, then washed again in PBS-Tween. Slides were incubated with Tyr-Cy3 reagent (1:50) (Perkin Elmer) for 5 minutes, then washed in deionized water, stained with DAPI (Thermo Fisher Scientific), and mounted for image acquisition. All images were captured using a confocal microscope (Leica SP8) and LAS X software (Leica). Images were analyzed with Imaris software (Oxford Instruments). FoxO1 mean intensity of fluorescent (MFI) in cytoplasm was divided by nuclear FoxO1 MFI of the same cell to determine ratio cytoplasmic/nuclear, lower ratio indicated increased nuclear localization.

### scRNA-seq and analysis

2,000 sgCtrl or sgIcos P14 cells were transferred into 3 recipient mice each before infection with LCMV cl13 (and CD4 depletion). Five weeks post infection, spleens were collected and single cell suspension from each mouse was surface stained with CD44 (FITC), CD8 (PerCP), PD-1 (PE-Dazzle), CD45.1 (APC) and live/dead IR for 30 min on ice. Samples were washed 3 times with PBS, 1%EDTA, 2% FBS. Then each single cell suspension was divided in two and stained with a unique cell multiplex oligonucleotide tag (CMO) from the 3’ CellPlex Kit set A (10x Genomics) and sort separately. P14 CD8 T cells were sorted as Live CD8^+^ CD45.1^+^ by FACS (BD Aria). Viability of the sorted cells was assessed using Acridine Orange/Propidium Iodide viability staining reagent (Nexcelom), showing a viability of 97%. Cells from all mice were then pooled in equal proportions.

scRNA-seq was performed on the Chromium microfluidics platform (10x Genomics) using the 3’ gene expression (3’ GEX) V3.1 kit, targeting a recovery of 35,000 cells per lane. Briefly, Gel-Bead in Emulsions (GEMs) were generated on the sample chip in the Chromium X system. Barcoded cDNA was extracted from the GEMs after Post-GEM RT-cleanup and amplified for 11 cycles. Full-length cDNA from poly-A mRNA transcripts underwent enzymatic fragmentation and size selection to optimize cDNA amplicon size (∼400 bp) for library construction. The cDNA fragments were then subjected to end-repair, adapter ligation, and 10x-specific sample indexing according to the manufacturer’s protocol (10x Genomics). The concentration of the single-cell library was accurately quantified using qPCR (Kapa Biosystems) to achieve appropriate cluster counts for paired-end sequencing on the NovaSeq 6000 (Illumina), aiming for a sequencing depth of 25,000 reads pairs per cell for gene expression libraries and 2,000 reads pairs per cell for CMO libraries. Raw fastq files were processed with the mm10 reference genome (2020-A) and demultiplexed using Cell Ranger v7.0.1 (10x Genomics).

Downstream analysis was performed in R v4.3.2 using Seurat v5 package^84^. Cells with greater than 5% mitochondrial transcript unique molecular identifier (UMI) counts, greater than 3000 total transcript UMI count, or less than 1100 transcript UMI count were excluded. In total, we retained 12,654 cells for analysis. Standard analysis was run: SCTransform RunPCA to identify the top 30 principal components, RunUMAP, FindNeighbors, FindClusters. Cell cycle scores (calculated using the CellCycleScoring function in Seurat) and mitochondrial percentage were selected as variables to regress out during SCTransform. Samples were integrated with 5000 variable genes chosen for the anchoring process. Clustering was visualized using Clustree^85^, and a cluster resolution of .09 was chosen for five main clusters. Tpex and effector-like cells were subset and re-clustered with the same pipeline as described above and clustered with a resolution of .17 and .09, respectively. SgIcos and sgCtrl samples were pseudobulked using function Seurat2PB. Differentially expressed genes were determined with edgeR^86^ using a generalized linear model quasi-likelihood F test and defined as having FDR < 0.05 and log2(FC) > 0.5. To annotate the defining genes of each cell cluster, we checked for the presence of well-characterized marker genes for T cells within the differentially expressed genes of a given cluster, and assigned each cluster an identity. Gene set scoring for single cell analysis was performed using VISION R package 3.0.1, uMAP plots show the top quintile of cells enriched for the signature highlighted in blue. Gene set was used from Giles *et al.* (Tex-KLR)^44^.

### Tumor burden assessment by luciferase detection

In vivo bioluminescence imaging was performed using an IVIS Spectrum system (Caliper LifeSciences) to quantify liver tumor burden before being evenly assigned to various treatment study cohorts. Mice were imaged 5 minutes after intraperitoneal injection with D-luciferin (150 mg/kg) (Thermo Scientific). Luciferase signal was quantified using Living Image software (Caliper LifeSciences). Normalized luciferase signal was calculated by subtracting the background signal. We ensured each treatment cohort had equivalent average luciferase signal before treatment initiation. Mice with low luciferase signal (10-fold lower than average signal) 5-7 days after hydrodynamic injection were excluded from the study.

### Tumor Infiltrating Lymphocyte (TIL) isolation

Liver was weighed after collection and approximately 1g was used for further processing. Liver was minced into pieces and digested in 15ml of RPMI (10%FBS) with collagenase IV (C5138, Sigma) 0.25mg/ml and DNaseI (DN25, Sigma) 0.1mg/ml 30min at 37°C. Further homogenization of tissue was performed with 10ml syringe and 16G needles, then filtered through 100µm strainer. To enrich for immune cells, cells from liver were resuspended in 25% Percoll (Citiva/Fisher) and underlaid with 70% Percoll (Citiva/Fisher), spun at 400g for 20min without acceleration and brake. Interface containing immune cells was collected and subjected to ACK lysis buffer (Gibco) for 3min at RT to lyse remaining red blood cells.

---

**Fig. S1:**
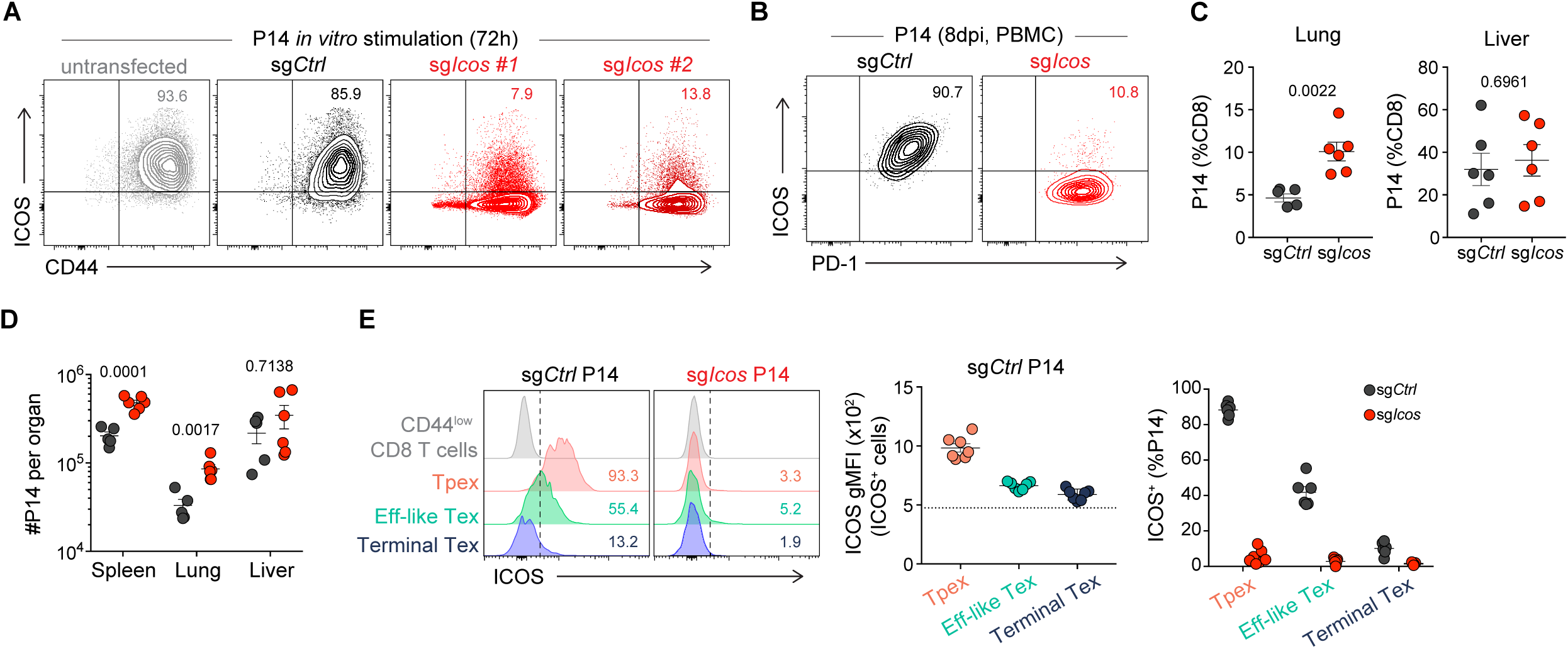
Experimental layout as in Fig. 1A. **(A)** Contour plots show CD44 and ICOS expression on P14 stimulated *in vitro* for three days with CD3/CD28 beads. P14 were untransfected (grey) or electroporated before culture with Cas9/sgCtrl (black), or two different sgRNA targeting Icos (red). **(B)** Contour plots show ICOS expression on sgCtrl and sgIcos P14 from blood of recipient mice, 8 days post LCMV infection. P14 frequency **(C)** and absolute number **(D)** in lung and liver of chronically infected animals. **(E)** Representative histograms show ICOS expression on progenitor, effector-like Tex, and terminal Tex P14 sgCtrl and sgIcos. Graphs show ICOS level of expression (geometric mean fluorescence intensity, gMFI, middle) and ICOS^+^ frequency among each subset (right). Data are representative of 3 independent experiments with five to six mice per group. Error bars show SEM. (C-D) unpaired Student’s t-test.

**Fig. S2:**
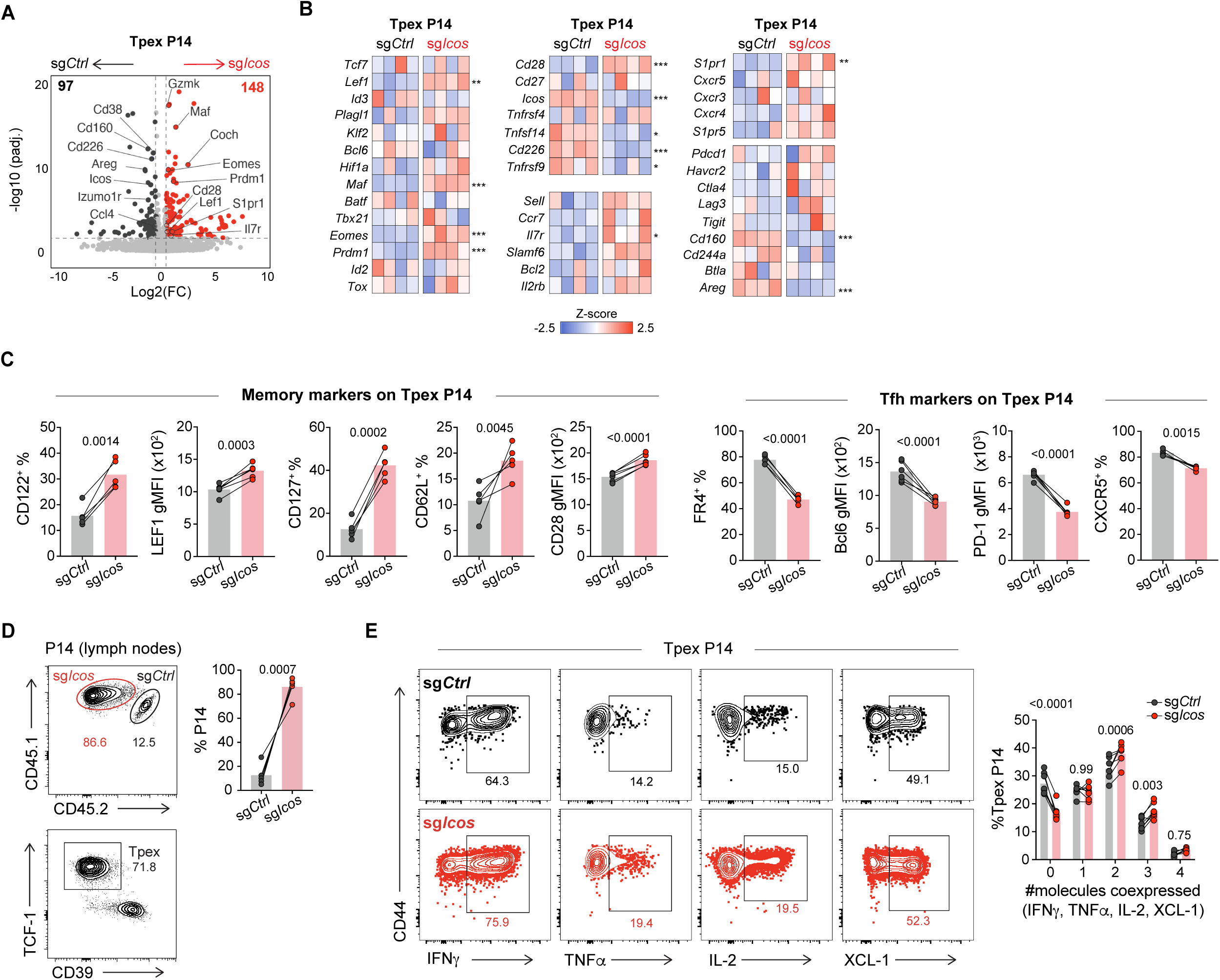
**(A-B)** RNAseq analysis of sorted control and sgIcos Tpex P14. **(A)** Volcano plot shows differentially expressed genes. Genes downregulated in ICOS-deficient Tpex are represented with black symbols, and genes upregulated in sgIcos Tpex are represented with red symbols to the right. Vertical dashed lines indicate 1.5-fold change; horizontal dashed lines indicate adjusted P-value=0.05. **(B)** Heatmap show relative expression of key genes in control and ICOS-deficient Tpex P14. **(C)** Expression of memory T cell markers (CD122, LEF1, CD127, CD62L, CD28) and Tfh markers (FR4, Bcl6, PD-1, CXCR5) on co-transferred sgCtrl and sgIcos Tpex P14 from spleens of chronically infected animals. **(D)** Frequency of co-transferred P14 in lymph nodes (axillary, brachial, inguinal and cervical), and frequency of Tpex among P14. **(E)** Contour plots show frequency of cytokine producing cells (IFNγ, TNFα, IL-2 or XCL-1) among splenic Tpex sgCtrl and sgIcos P14 (35 DPI). Graph show number of cytokines/chemokines co-produced by control or sgIcos Tpex P14. Data in (C-D) are representative of 3 independent experiments with five to six mice per group. Symbols represent individual mice with bar showing the mean value of all animals analyzed. Connecting lines between symbols indicate cells analyzed from the same mouse. (B) Z-score was calculated as (sample TPM - mean of all TPM) divided by standard deviation. Significance was determined by Independent hypothesis weighting (IHW) on p-values from DESeq2 analysis. (C-D) paired Student’s t-test; (E) two-way ANOVA with Sidak’s correction for multiple comparisons.

**Fig. S3:**
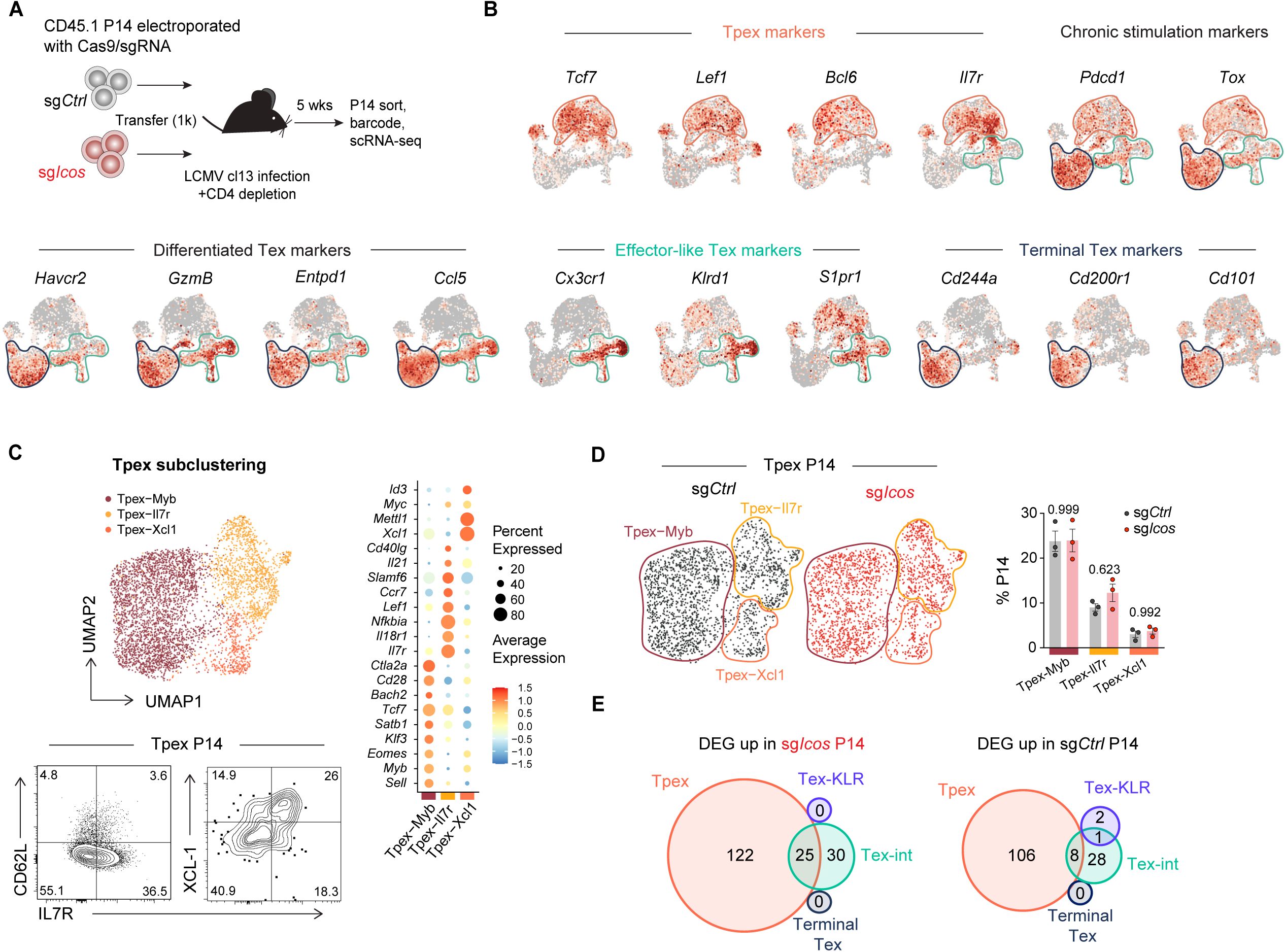
**(A)** sgCtrl and sgIcos P14 were transfered into separate recipient mice before LCMV cl13 infection (helpless). Five weeks post infection, P14 were sorted, barcoded and pooled for scRNA-seq. **(B)** UMAP of selected genes. **(C)** UMAP of Tpex P14 subclustered (top panel). Contour plots show expression of CD62L, IL7R and XCL-1 by Tpex P14 (bottom panel). Dot plot representing average and percent expression of selected genes (right panel). **(D)** UMAP of control (black) and ICOS-deficient (red) Tpex P14, graph of relative distribution of clusters. **(E)** Venn diagram comparing differentially expressed genes in control and ICOS-deficient Tpex, Tex-Int, Tex-KLR and terminal Tex. (D) Symbols represent individual mice with bar showing the mean value of all animals analyzed and bars indicate SEM. (D) Significance was determined using one-way ANOVA with Holm-Sidak’s correction for multiple comparisons.

**Fig. S4:**
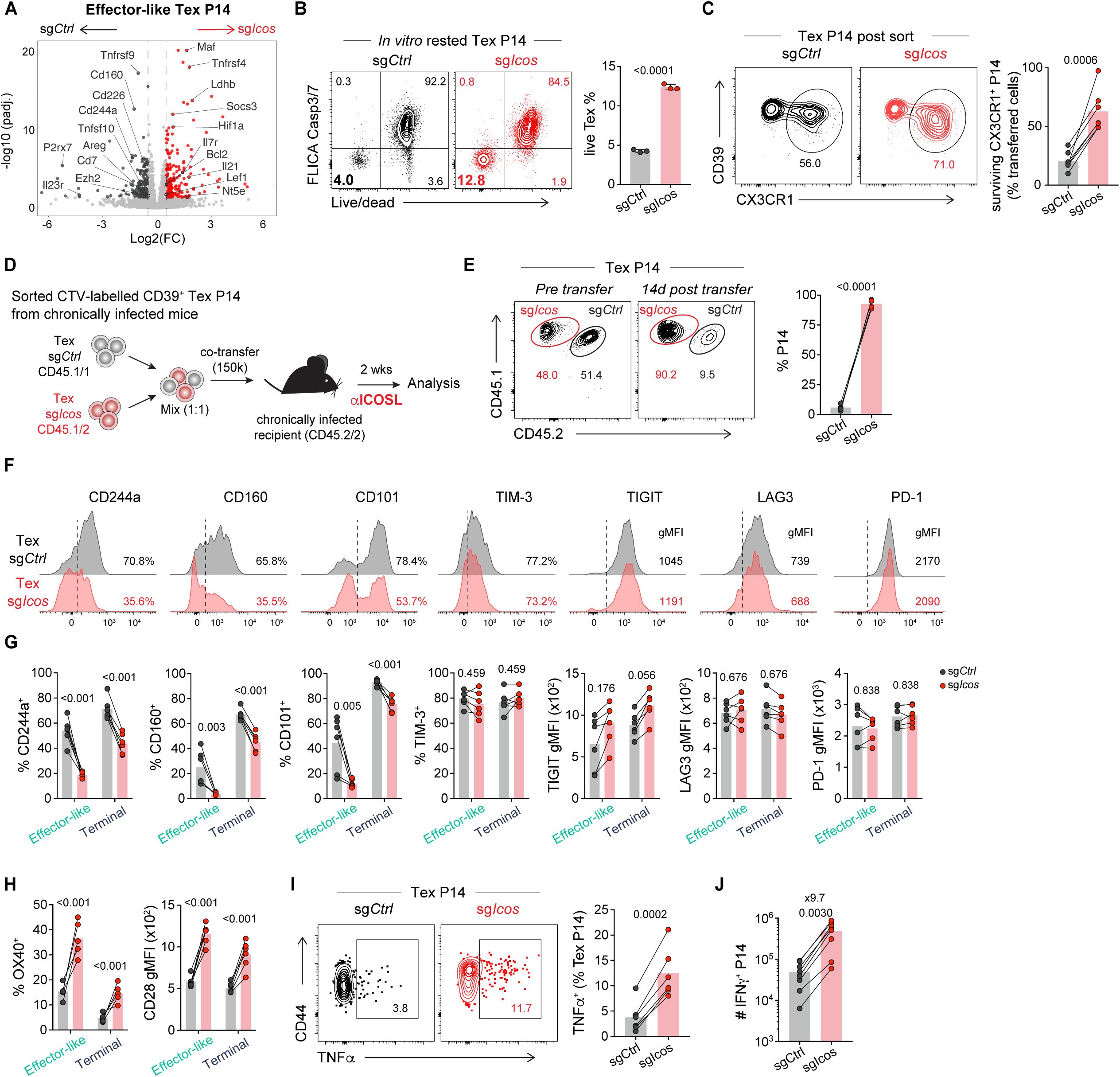
**(A)** Volcano plot showing differentially expressed genes between control and ICOS-deficient effector-like Tex P14 based on RNA sequencing of respective sorted P14. Genes downregulated in ICOS^-^deficient cells are shown in black symbols, and genes upregulated are shown in red. Vertical dashed lines indicate 1.5-fold change; horizontal dashed lines indicate adjusted P-value=0.05. **(B)** Contour plots and graph show frequency of live cells among sorted sgCtrl and sgIcos Tex (CD39^+^) rested *in vitro* for 3d without stimulation. **(C)** Contour plots show frequency of CX3CR1^+^ effector-like Tex among control and sgIcos P14 pre-transfer, and graph shows frequency of surviving effector-like Tex P14 in spleen two weeks post transfer, as percentage of original number of CX3CR1^+^ Tex P14 adoptively transfer (estimating 10% engraftment). **(D)** Experimental layout, similar to Fig 4F, but mice received αICOSL after adoptive Tex transfer. **(E)** Contour plots and graph show relative frequencies of control and sgIcos Tex pre and 2 weeks post transfer into chronically infected recipients. **(F)** Representative histograms and **(G)** graphs show inhibitory receptors expression on control (black) and ICOS-deficient (red) Tex (CD39^+^) P14. **(H)** Expression of costimulatory molecules CD28 and OX-40 on control and ICOS-deficient effector-like (CX3CR1^+^) and terminal (CX3CR1^neg^) Tex P14. **(I)** Contour plots and graph show frequency of TNFα-producing cells among effector-like and terminal Tex P14 after GP33 peptide restimulation. **(J)** Absolute number of IFNγ-producing P14 sgCtrl and sgIcos in the spleen of chronically infected animals. Data in (F-J) are representative from 3 independent experiments with five to six mice per group. Data in (B) are representative from 2 independent experiments with three samples per group. Data in (E) are from one experiment with four mice per group. (B) mean +/- SEM. (F-J) Connecting lines between symbols link cells analyzed from the same mouse, with bar showing the mean value. (F) Vertical dashed line indicate positivity. (B) unpaired Student’s t-test, (E,G-J) paired Student’s t-test

**Fig. S5:**
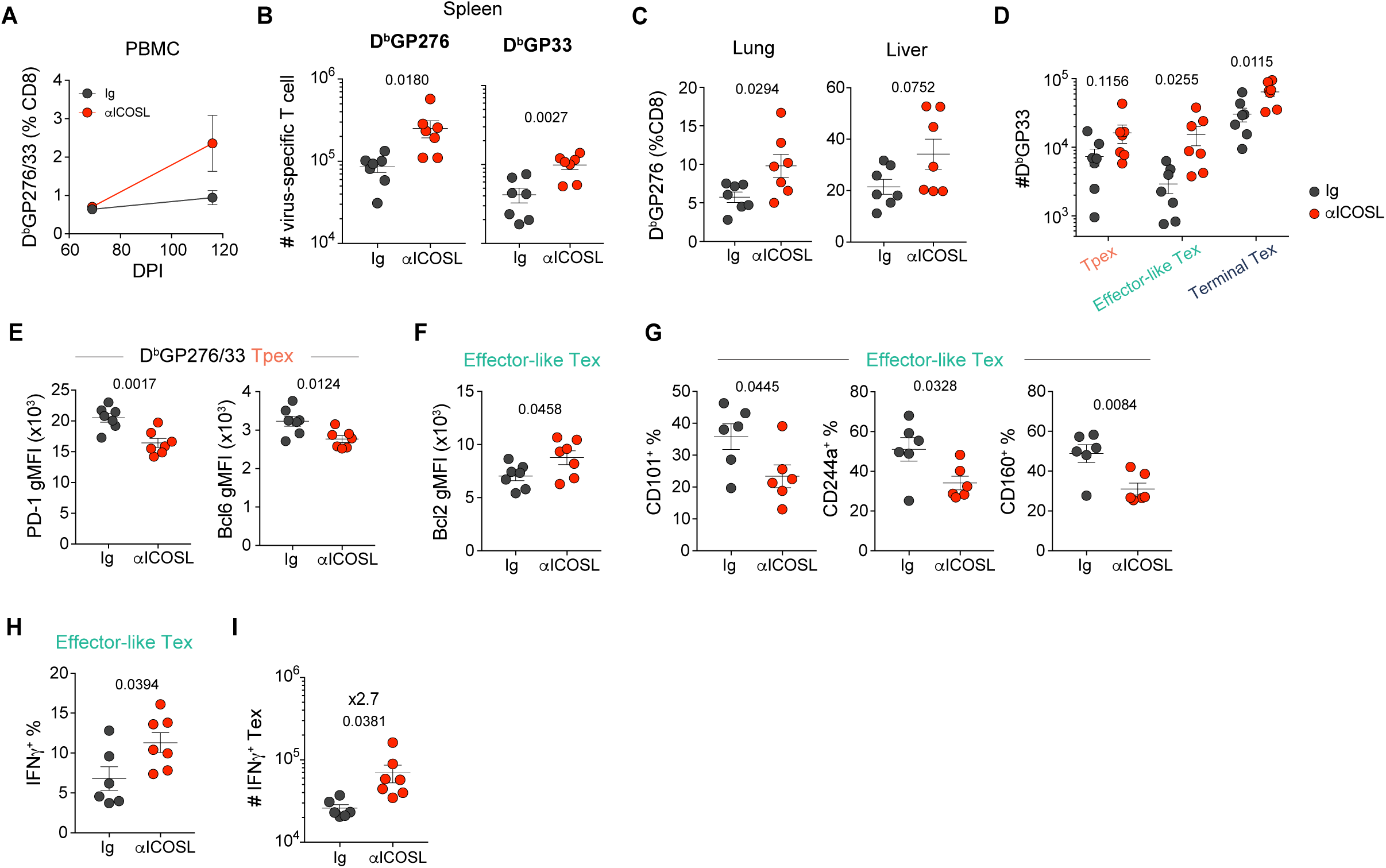
Experimental layout as in Fig.5A. **(A)** Frequency of LCMV-specific CD8 T cells in the blood pre and post treatment. **(B)** Absolute number of splenic LCMV-specific CD8 T cells. **(C)** Frequency of LCMV-specific CD8 T cells (D^b^GP276) in lung and liver. **(D)** Absolute number of TCF1^+^ progenitor, CX3CR1^+^ effector-like, and CX3CR1^neg^TCF-1^neg^ terminal Tex LCMV-specific (D^b^GP33) CD8 T cells. **(E)** gMFI of PD-1 and Bcl6 of LCMV-specific Tpex. **(F)** Bcl2 expression on virus-specific effector-like Tex. **(G)** Expression of inhibitory receptors (CD101, CD244a, CD160) on virus-specific effector-like Tex. **(H)** IFNγ production by CX3CR1^+^ PD-1^+^ CD8 T cells after LCMV-peptides restimulation. **(I)** Absolute number of IFNγ^+^ CD39^+^ PD-1^+^ T cells in the spleens of chronically infected animals. Data in (A-I) are representative from 2 independent experiments with five to seven mice per group. Symbols represent individual mice. Data are shown as mean +/- SEM. (A-I) unpaired Student’s t-test.

**Fig. S6:**
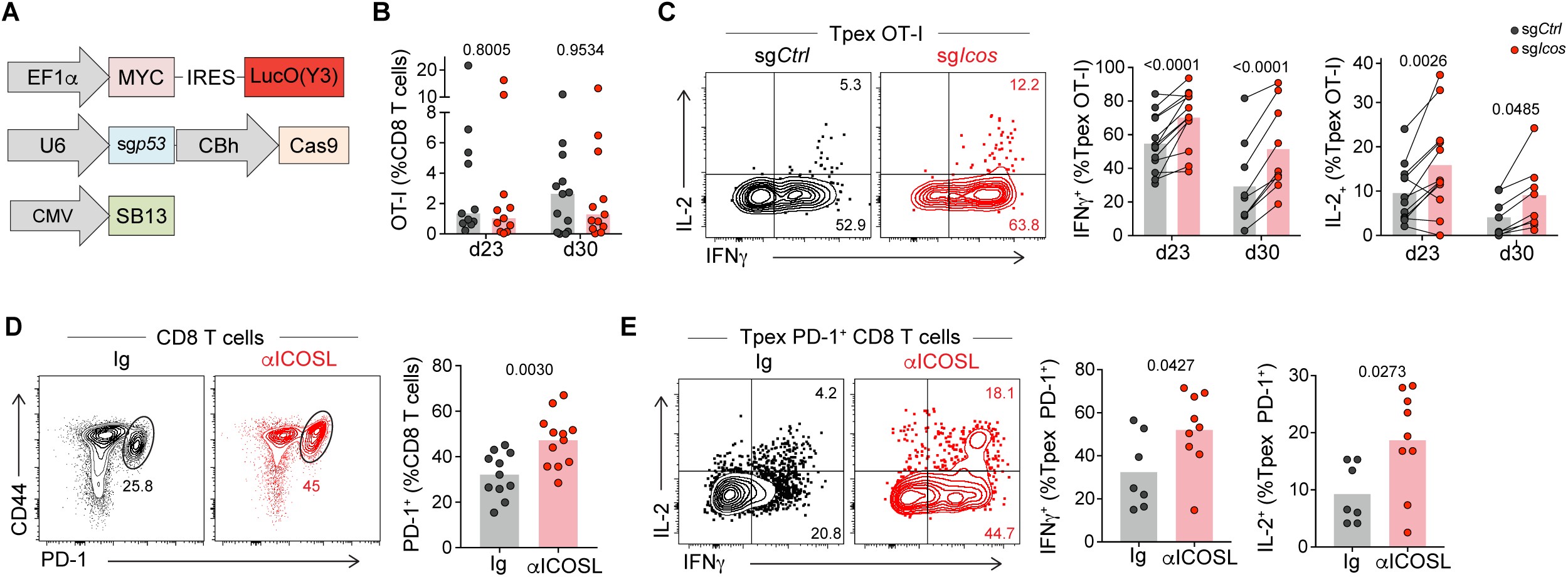
**(A)** Schematic of plasmids hydrodynamically injected into mice to induce HCC formation **(B-C)** Related to layout Fig.6D. **(B)** Frequencies of co-transferred OT-I at d23 and d30 post tumor initiation. **(C)** Frequency of IFNγ and IL-2 producing Tpex OT-I. **(D-E)** Related to layout Fig.6H. **(D)** Frequency of PD-1^+^ CD8 T cells. **(E)** Frequency of IFNγ and IL-2 producing Tpex PD-1^+^ CD8 T cells. Data in (B-E) are representative from 2 independent experiments with five to eight mice per group. Symbol represent individual mice. Connecting lines between symbols link cells analyzed from the same mouse. Data are shown as mean. (B-C) paired Student’s t-test; (D-E) unpaired Student’s t-test.

